# A Cdh3-Lam332 signaling axis in a leader cell subpopulation controls protrusion dynamics and tumor organoid collective migration

**DOI:** 10.1101/2022.05.10.491382

**Authors:** Priscilla Y Hwang, Jairaj Mathur, Yanyang Cao, Jose Almeida, Daphne Cornish, Maria Clarke, Amit Pathak, Gregory D Longmore

## Abstract

Carcinoma dissemination can occur when heterogeneous tumor and tumor stromal cells clusters migrate together via collective migration. Cells at the front lead and direct collective migration, yet how these leader cells form and interact with the microenvironment to direct migration are not fully appreciated. From live videos of primary mouse and human breast tumor organoids in a 3D microfluidic system that mimics the native breast tumor microenvironment, we developed 3D computational models which hypothesize that leader cells generate high protrusive forces and overcome extracellular matrix (ECM) resistance. Using single cell sequencing, we reveal leader cells are heterogeneous, and identify and isolate a unique Cadherin-3 (Cdh3) positive leader cell subpopulation that is necessary and sufficient to lead migration. Cdh3 controls leader cell protrusion dynamics through the local production of Laminin-332 which is required for integrin/focal adhesion function. Our findings highlight how a subset of leader cells interact with the microenvironment to direct collective migration.

**Teaser:** Higher protrusions of Cdh3+ leader cells polarize tumor organoids that then invade collagen via Lam332 adhesion feedback.

## Introduction

Collective migration is the process by which multiple cells move in a coordinated manner and is essential for many physiological processes during development and wound healing (1-3). In disease states, such as cancer, collective migration is also important for tumor invasion and migration that lead to metastases(4, 5). Unlike single cell migration, collective migration requires that cells interact with neighboring cells as well as respond to environmental cues(1, 6, 7). Collectively migrating cells can be broadly viewed as consisting of two general populations: leader cells and follower cells(8-11). Leader cells locate to the leading edge in the direction of migration while follower cells are at the rear(10). Many studies have characterized the cells at the leading edge, and it is now appreciated that leader cells explore their immediate microenvironment (chemical and physical), determine the direction of migration, generate traction forces necessary to move the whole, respond to and remodel their structural environment to facilitate invasion, and finally transmit signals to follower cells(9, 12). Despite this, the full repertoire of molecular mechanisms that drive leader cell formation and function and in what contexts is not fully appreciated.

Within the extracellular matrix (ECM) surrounding tumors and in the circulating blood stream, tumor cells can be found in clusters and migrate through the blood and exit the bloodstream (extravasate) as clusters. Blood tumor clusters have up to a 100-fold greater metastatic potential than single-cell circulating tumor cells(13-16). In breast cancer, leader cells(8, 10) are identified by the expression of the intermediate filament protein cytokeratin 14 (K14)(17-19) and in spontaneous mouse models of breast cancer, multicellular tumor clusters that contain K14+ cells have higher propensity to metastasize(19, 20).

One hypothesis for tumor leader cell development posits that the cells at the cluster edges, in contact with ECM, become leader cells in response to local microenvironmental cues(12, 21). However, in other study systems, K14+ cells within primary breast tumor organoids, can be randomly distributed and in the presence of hypoxia and exposure to chemical or mechanical signaling gradients migrate within the cluster to the leading edge, without any apparent change in cell fate or number(22). These observations suggest that so-defined leader cells could be more heterogeneous than previously appreciated, exhibit complex responses to environmental signals, and behave distinctively from follower cells.

Many studies of collective invasion, including tumor invasion, make use of aggregated cell lines, not primary tissues or tumors(23, 24). While extremely useful for identifying cell biological processes and important molecules and their function(s) therein, collectively migrating primary tumors, or primary cells in general, are inherently more heterogeneous in their cellular composition and behavior(25, 26). Moreover, potently metastatic tumor cell clusters contain not only leader and follower tumor cells, but also stromal cells such as cancer-associated fibroblasts (CAFs) and immune cells(14, 25-28), and when stromal cells are present metastatic potential is increased(29, 30). At present we do not have a comprehensive understanding of the cellular complexity within invasive primary tumor clusters, the functions of the different cell types within the clusters, nor the contribution of various environmental signals to the behavior of different cell types within the tumor cluster to collective migration.

Experiments studying collective invasion, *ex vivo*, typically expose cells to uniform environmental signals and ambient oxygen concentration, which do not reflect the *in vivo* exposure of invasive tumor cell clusters(31). Indeed, the cellular, chemical, and physical tumor microenvironment is extremely dynamic(32-34). Therefore, these studies in non-pathophysiologic environmental conditions are limited in their biologic implications and clinical translational potential. To address these limitations, efforts investigating leader cell driven collective migration must take into consideration cellular and environmental signal heterogeneity as well as signaling gradients.

To address these limitations, we made use of an *ex vivo* 3D microfluidic system and genetically defined primary breast tumor organoids derived from a metastatic genetically engineered mouse model (GEMM) and a human metastatic triple negative breast cancer (TNBC) PDX (human in mouse) model as these incorporate native tumor cell heterogeneity as well as tumor associated stromal cells. Tumor organoids are exposed to pathophysiologic hypoxia (< 2%) and various environmental signaling gradients within a 3D collagen I matrix(22). To gain insights into how leader cells polarize to the leading edge and then lead directed collective migration in 3D, we performed computational modeling or simulations based upon observations from published live cell 3D videos(22). To determine the extent of cellular heterogeneity of leader cells and gain molecular and signaling pathway insights during active collective migration we performed single cell RNA sequencing (scRNAseq) molecular analysis of collectively migrating primary tumor clusters in response to different environmental signals. Informed by results from scRNAseq analysis we were able to experimentally test hypotheses generated from precedent computational modeling.

We identify significant K14+ “leader” cell heterogeneity that differs depending upon the environmental signal. We isolate a subset of K14+ tumor cells and show that they are necessary and sufficient to lead collective migration of non-migratory normal breast organoids. These leader cells are enriched for expression of Cdh3 and Cdh3 is required for leader cell polarization and function. In leader cells, Cdh3 controls protrusion dynamics and overcomes ECM resistance to migration by controlling β-catenin/TCF regulated transcription and local production of the basement membrane component Laminin 332 which is required for collective migration by controlling Integrin/focal adhesion (FA) function, and thus, protrusion dynamics in leader cells.

## Results

### Computational Modeling of tumor organoid collective invasion in 3D predicts for leader cells defined by high protrusive forces and ECM mediated feedback

As tumors progress to invade and metastasize leader cells that direct collective invasion arise, yet how they form and function to direct collective invasion/migration of tumors are not fully understood. In previous work we designed and characterized a microfluidic device to study directed collective migration of primary mouse or human breast tumor organoids through a 3D collagen I matrix under hypoxic conditions that are present in primary tumors (< 2% O_2_), and in response to either a chemical signaling gradient (SDF1) or a mechanical signaling gradient (interstitial flow)(22). In this system, K14+ breast tumor cells (leader cells) are initially randomly distributed throughout the tumor organoid and comprise between 10-20% of total cells. In contrast to other systems of breast tumor organoid invasion(19), K14+ cells are not solely present at the organoid surface where tumor cells contact the collagen I matrix. When tumor organoids are exposed to various environmental signaling gradients K14+ cells first polarize to what will become the leading edge and then lead “spherical” collective migration up an SDF1 chemical gradient or with interstitial fluid flow gradient.

To gain insight and generate experimentally testable hypothesis into how leader cells polarize to the leading edge and lead directed collective migration, we developed theoretical and computational models of tumor organoid migration in 3D. Simulation of whole organoid movement were calibrated against videos of GFP-labeled leader cells in WT mouse PyMT breast tumor organoids in hypoxia (< 2%) migrating through Collagen I matrices in response to an SDF1 chemical gradient (22). The key goal of this modeling effort was to understand and predict adhesions and forces that leader cells must follow to generate the directed collective organoid movement as observed in videos.

We used a lattice-based modeling approach with energy functionals to define different cell types, adhesions, and protrusive forces (Fig. 1b). In spatiotemporal simulations, stochastic movements are determined based on an energy-minimization criterion calculated over defined Monte Carlo steps (details in Methods). From 10-20% K14+ leader cells observed in primary tumor organoids, we modeled a 3D tumor organoid with 10% leader cells (green), a cell population whose properties we will study, mixed randomly with non-leader “follower” cells (blue) and a rightward chemokine (SDF1-like) gradient (Fig. 1a,b). From live movies of Actin.GFP-labelled K14+ cells in tumor organoids, in the microfluidic device, that collectively migrated in 3D collagen I,(22) we observed three key phenotypes: (1) leader cell polarization to the migratory front, calculated in simulations as percentage of leader cells in the front 60° cone relative to all leader cells; (2) organoids stay intact and retain circular shape, defined here as circularity (minor axis divided by major axis of a fitted ellipsoid); and (3) net organoid displacement towards the direction of chemokine gradient. In our simulations, we calculated and plotted these three readouts to calibrate against video observation of primary breast tumor organoid collective migration in 3D.

**Figure 1.**
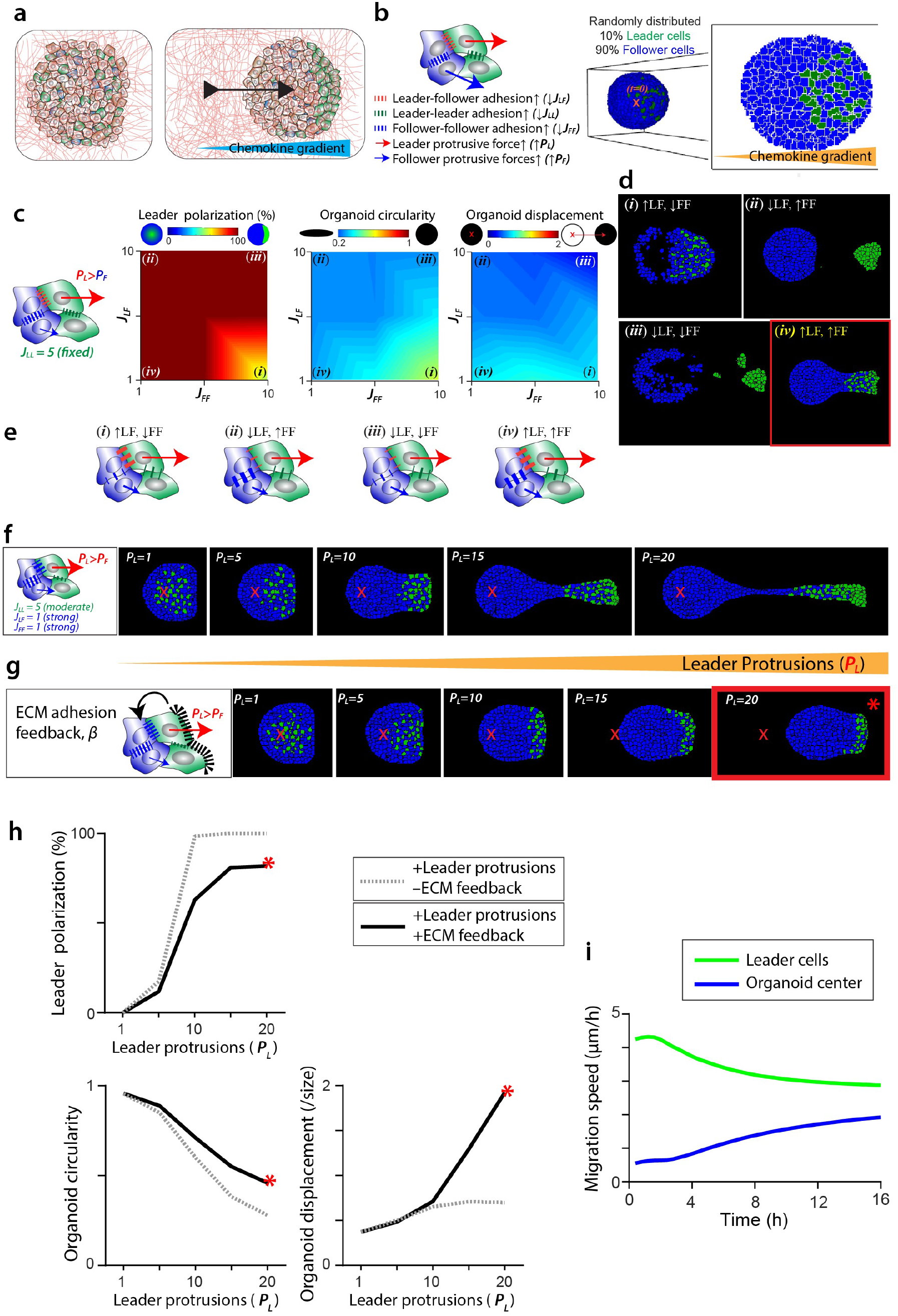
Computational Modeling of tumor organoid collective invasion in 3D. **(a)** Schematic describing our experimental observations of how a 3D tumor organoid with heterogeneous cell population responds to a chemokine gradient (SDF1) with leader cell polarization and whole organoid displacement along the gradient. **(b)** Key modeling variables of differential leader/follower adhesions and protrusions (left panel) implemented in a modified cellular Potts model with mixed leader and follower population of tunable properties. Right panel illustrates the position of leaders and followers in 3D at time *t*, and a slice view through the middle of the organoid, with rightward chemokine gradient. Left panel shows a schematic of a leader cell (green), the surrounding follower cells (blue), leader/follower protrusions (*P*_*l*_ *and P*_*F*_) along with leader-follower adhesions (*J*_*LF*_, red), leader-leader adhesions (*J*_*LL*_*a* green) and follower-follower adhesions (*J*_*FF*_, blue). **(c)** Heatmaps for leader cell polarization (left), organoid circularity (middle) and organoid displacement relative to its initial radius (right) for varying *J*_*LF*_ and *J*_*FF*_, with moderate leader-leader adhesions (*J*_*LL*_ = 5) and high leader cell protrusive forces (*P*_*L*_ = 20, *P*_*F*_ = 1). For *J*_*LL*_ = 1,10 refer to suppl. Fig. S1 b,c. **(d)** Cross-section views of 3D organoid configuration at *t* = 4 *hours* for high leader cell protrusive forces with adhesions cases: (*i*) weak follower-follower adhesions (*J*_*FF*_ = 10) and strong leader-follower adhesions (*J*_*LF*_ = 1); (*ii*) strong follower-follower adhesions (*J*_*FF*_ = 1) and weak leader-follower adhesions (*J*_*LF*_ = 1); (*iii*) weak follower-follower adhesions (*J*_*FF*_ = 10) and weak leader-follower adhesions (*J*_*LF*_ = 10); (*iv*) strong follower-follower adhesions (*J*_*FF*_ = 10) and strong leader-follower adhesions (*J*_*LF*_ = 10). **(e)** Schematics describing the above 4 cases. **(f)** Cross-section views of 3D organoid configuration at *t* = 16 *hours* for increasing *P*_*L*_ = 1 *to* 20. Here, ‘x’ symbol (in red) marks the initial position of organoid center. **(g)** With ECM adhesion feedback present, cross-section views of 3D organoid configuration at *t* = 16 *hours* for increasing *P*_*L*_ = 1 *to* 20. Red outline and ‘*∗*’ symbol (in red) marks the case which captures the experimental observations. **(h)** For the conditions without ECM adhesion feedback (black dotted line) and with ECM adhesion feedback (black solid line), leader cell polarization, organoid circularity, and organoid displacement relative to organoid radius are plotted for varying leader protrusions *P*_*L*_ = 1 − 20. **(i)** Average leader population speed (green line) and average follower population speed (blue line) over time.

First, we tested whether differential adhesions between leaders and followers alone could explain leader cell polarization to the front and dragging the entire organoid with them. We systematically varied leader-follower (*J*_*LL*_), follower-follower (*J*_*FF*_), and leader-leader adhesions (*J*_*LL*_), while keeping leader and follower protrusions the same *P*_*L*_ = *P*_*F*_ = 1. We found that there was no combination of cell-cell adhesions that alone could generate leader cell polarization (Suppl. Fig. S1a; Suppl. Movie 1). Thus, differential cell-cell adhesions alone did not capture the ability of leader cells to polarize to the organoid front or enable organoid movement. Next, we hypothesized that for leader cells to move through the organoid they need to generate higher protrusive forces compared to follower cells, which we implemented by assigning higher values of leader cell protrusive force (*P*_*L*_ = 20; calibrated ahead). With this added variable, we repeated simulations for all combinations of cell-cell adhesions (Fig. 1c; Suppl. Fig. S1b,c). Indeed, with higher relative protrusive force, leader cell polarization increased for all values of *J*_*LL*_, *J*_*FF*_, and *J*_*LF*_ (Fig. 1c; Suppl. Fig. S1b,c). Based on these simulations, we concluded that higher protrusive forces were important for the leader cells to move through the organoid and polarize to the invasive front, as observed in live videos. Additionally, we calculated organoid circularity and organoid centroid displacement normalized by organoid radius after 16 hours (Fig. 1c; Suppl. Fig. S1b,c). For most combinations of cell adhesions, organoid circularity and its displacement remained low (Fig. 1c; Suppl. Fig. S1b,c), i.e., organoids did not move or stay intact as leader cells polarize.

Next, we fixed leader-leader adhesion strength as moderate, *J*_*LL*_ = 5, and analyzed organoid configuration after 4 hours for 4 cases of leader-follower and follower-follower adhesions (Fig. 1d,e; Suppl. Movies 2-5). Case (*i*): strong leader-follower (*J*_*LF*_ = 1) and weak follower-follower (*J*_*FF*_ = 10) adhesions made it more energetically favorable to maintain leaders with followers while followers scatter, which reduced leader cell polarization and broke up the organoid (Fig. 1c, d(i); Suppl. Movie 2). Case (*ii*): weak leader-follower (*J*_*LF*_ = 10) and strong follower-follower (*J*_*FF*_ = 1) adhesions reduced organoid scattering, but leader cells exited the organoid after frontward polarization (Fig. 1c, d(ii); Suppl. Movie 3). Case *(iii)*: both leader-follower and follower-follower weak adhesions (*J*_*FF*_ = *J*_*LF*_ = 10) allowed for leader cell polarization but the organoid dissociated and leader cells exited the organoid (Fig. 1c, d(iii); Suppl. Movie 4). Thus, if either leader-follower or follower-follower adhesions (*J*_*LF*_*a J*_*FF*_) were weak, the organoid integrity was compromised. In Case (*iv*), upon setting strong leader-follower and follower-follower adhesions (*J*_*LF*_ = *J*_*FF*_ = 1), the organoid remained intact, leader cells polarized to the front, and they did not exit the organoid (Fig. 1c, d(iv); Suppl. Movie 5), which captured organoid morphology, yet organoid displacement remained low. Based on these parametric scans, we set moderate leader-leader adhesions (*J*_*LL*_ = 5) and strong follower-follower and leader-follower adhesions (*J*_*LF*_ = *J*_*FF*_ = 1). To analyze the effect of leader protrusive forces, we varied *P*_*L*_ from 1 to 20 and observed organoid configuration after 16 hours (Fig. 1f). With increasing leader protrusions, we predict increasing leader cell polarization, decreasing organoid circularity, and increasing organoid displacement (Fig. 1f, h). However, even for the highest value of *P*_*L*_ = *f*0, the organoid became highly elongated and predicted organoid movement was minimal relative to its size (Fig. 1f,h; Suppl. Movie 6).

Thus far, although preferential protrusions in leader cells moved them to the front of the organoid, this polarization had no functional advantage for the rest of the organoid. After leader cells arrive at the front, one key change in their state is the presence of the ECM, which is less inside the organoid. To understand possible functional advantages of leader cell polarization, we hypothesized that upon reaching the organoid front leader cells form adhesions with the surrounding ECM (collagen fibers) and that these events generate an active adhesion feedback for the follower cells. Such ECM adhesions and active feedback (implemented as parameter *β*; see model details in Methods) work against the ECM resistance to enable invasion through energy minimization in the model. With this ECM adhesion feedback, we performed simulations for varying leader protrusive forces, while keeping cell adhesions the same as established above (Fig. 1f). For *P*_*L*_ < 5, there was little change in the organoid configuration (Fig. 1g), leader cell polarization, circularity, and net organoid displacement (Fig. 1h). For higher values of leader protrusions (*P*_*L*_ > 5), one key advantage of the ECM feedback is that the organoid circularity is maintained even as the leader cell polarization increases. For *P*_*L*_ = 20 the organoid travels 2 times its radius in 16 hours, which matches video observations (Fig. 1g,h; Suppl. Movie 7). This final simulation (annotated by an asterisk in Figs. 1g,h) captured video observations per all three criteria of leader cell polarization, organoid circularity, and net organoid invasion, as noted above. For this condition of preferential leader protrusions along with an active ECM adhesion feedback, the average leader cell speed starts at a higher point relative to the rest of the organoid, and the speed of the whole organoid catches up as the leader cells reach the front (Fig. 1i), which was qualitatively consistent with live videos(22).

Based on this stepwise modeling approach of calibrating one parameter at a time, we make these following predictions for leader cell polarization and whole organoid invasion through 3D collagen: (1) differential cell-cell adhesions of leaders and followers alone do not explain leader polarization through organoid, (2) leader cells should have preferential protrusions relative to other cells (followers) in the organoid, and (3) at least a subpopulation leader cells forms active ECM adhesions upon arriving at the invasive front in order to reduce resistance presented by the collagen and thus direct the whole organoid movement. Experiments presented ahead will specifically test these predictions.

### Collectively migrating primary mouse PyMT breast tumor organoids exhibit K14+ “leader” cell heterogeneity

To gain insight into molecular regulation of K14+ “leader” cell function we performed single cell RNA sequence (scRNAseq) analysis of pooled mouse PyMT breast tumor organoids in hypoxia for 48h before exposure to a signaling gradient (no signal) and then after they had migrated collectively in response an SDF1 chemical gradient or Interstitial (IS) flow gradient (Fig. 2a-c, respectively; Suppl. Fig. S2).

**Figure 2.**
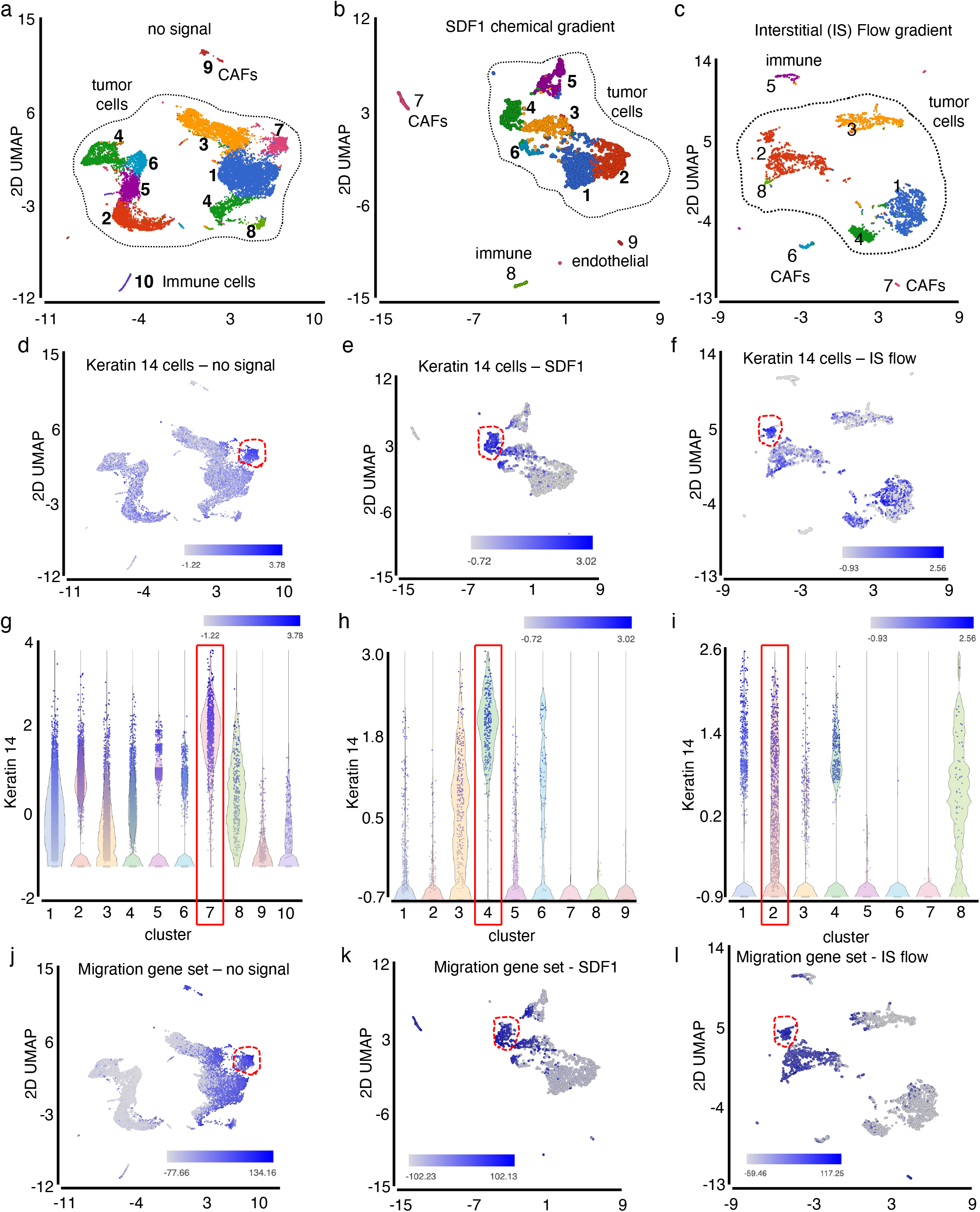
scRNASeq analysis: Collectively migrating primary mouse PyMT breast tumor organoids exhibit K14+ “leader” cell heterogeneity. **(a-c)** UMAP plot showing distinguished cell clusters for primary tumor organoids without migration (no signal), migration after SDF1 chemical gradient, and migration after interstitial (IS) flow gradient. **(d-f)** UMAP plot and **(g-i)** violin plots showing Keratin-14 (K14) expressing cell clusters for no signal, SDF1 chemical gradient, and interstitial (IS) flow gradient. **(j-l)** UMAP plot showing cell clusters expressing known migration gene sets.

scRNAseq analysis yielded data from 1500-2000 cells with a coverage of ∼125,000 read pairs per cell. To identify various cell types present we utilized graph-based clustering of transcripts, the ImmGen database, and published gene markers. Assessment of published cell-type markers resulted in elaboration of ten distinct cell clusters in organoids not exposed to signals, nine distinct cell clusters in response to the SDF1 gradient and eight distinct cell clusters in response to the IS flow gradient (Fig. 2 and Suppl. Fig. S2). There were nine, six, or five distinct tumor cell clusters, respectively, as defined by the presence of EpCAM and keratin 8, a small single cluster of endothelial cells in each (Pecam1+), and a single myeloid/immune cell cluster in each (CD45+). In each sample there were two distinct cancer associated fibroblast clusters: vascular CAFs (vCAFs) and matrix or mesenchymal CAFs (mCAFs)(35). Developmental CAFs (dCAFs)(35), if present, were interspersed within the tumor cell populations.

Prior to exposure to signals, K14 expression (a marker of leader cells in breast cancer) was distributed throughout all nine tumor cell clusters (Fig. 2d,g). In response to an SDF1 chemical gradient only three of six tumor cell populations were enriched for K14 expression (Fig. 2e,h), while in response to an IS flow gradient all five tumor cell populations expressed K14 (Fig. 2f,i). To increase resolution and better define the various K14+ sub-populations we probed the K14+ sub-populations in SDF1 treated organoids with known gene sets from GSEA. One cluster had high expression of cell proliferation markers (Suppl. Fig. S3a). All cell clusters were enriched for expression of hypoxia responsive genes (Suppl. Fig. S3b). Genes important for cell migration were expressed in all K14+ tumor cell clusters, as well as mCAFs and vCAFs (Fig. 2j-l). Interrogation of the data with a set of expressed genes enriched in K14+ tumor cells isolated from a primary mouse PyMT tumors (19) revealed that these genes were specifically enriched in tumor cell cluster seven of no signal, cluster four of SDF1 chemical gradient, or cluster two of the IS flow gradient (Fig. 2d-i, red outline; Suppl. Fig. S3c-e).

In sum, these molecular analyses indicated that there was significant K14 tumor cell molecular heterogeneity in collectively invading primary PyMT tumor organoids. Moreover, the K14+ leader cell heterogeneity appeared to vary depending upon the applied signal. In each of the samples, however, there appeared to be a unique cluster of K14+ tumor cells that exhibited genomic features associated with migration-capable leader cells.

### Cadherin 3 expression identifies a unique leader cell sub-population that is required for tumor collective migration

Are all K14+ tumor cell clusters capable of leading tumor collective migration? To address this question, we developed a method to isolate select K14+ tumor cell subpopulations and then developed a reconstitution assay using primary tumor cells to test if they could direct collective migration of normal, non-migratory, non-transformed primary mammary gland organoids.

We performed Volcano plots (Fig. 3a-c) comparing genes present in the migration capable, K14-enriched tumor cluster #7 of no signal, #4 of SDF1 chemical gradient, and #2 of IS flow gradient (Fig. 2g-l) with genes present in all other K14+ tumor cell clusters in each respective sample. A Venn diagram revealed 160 genes were common to all three subpopulations (Fig. 3d; Suppl. Table S1). To be able to isolate these K14+ subpopulations using FACS we asked which cell surface protein genes, known to be associated with tumor progression, were enriched in these 160 common genes. Examples included Cdh3, Jag2, Fgfr2, and Cntfr. Cadherin 3 (Cdh3) or P-Cadherin, in particular, was enriched in each (Fig. 3a-c; 3e-j). We confirmed the restricted expression of Cdh3 protein to K14+ leader cells by immunostaining migrating tumor organoids (Fig. 3k,l) and flow cytometric analysis of primary mouse PyMT breast tumors (Fig. 3m).

**Figure 3.**
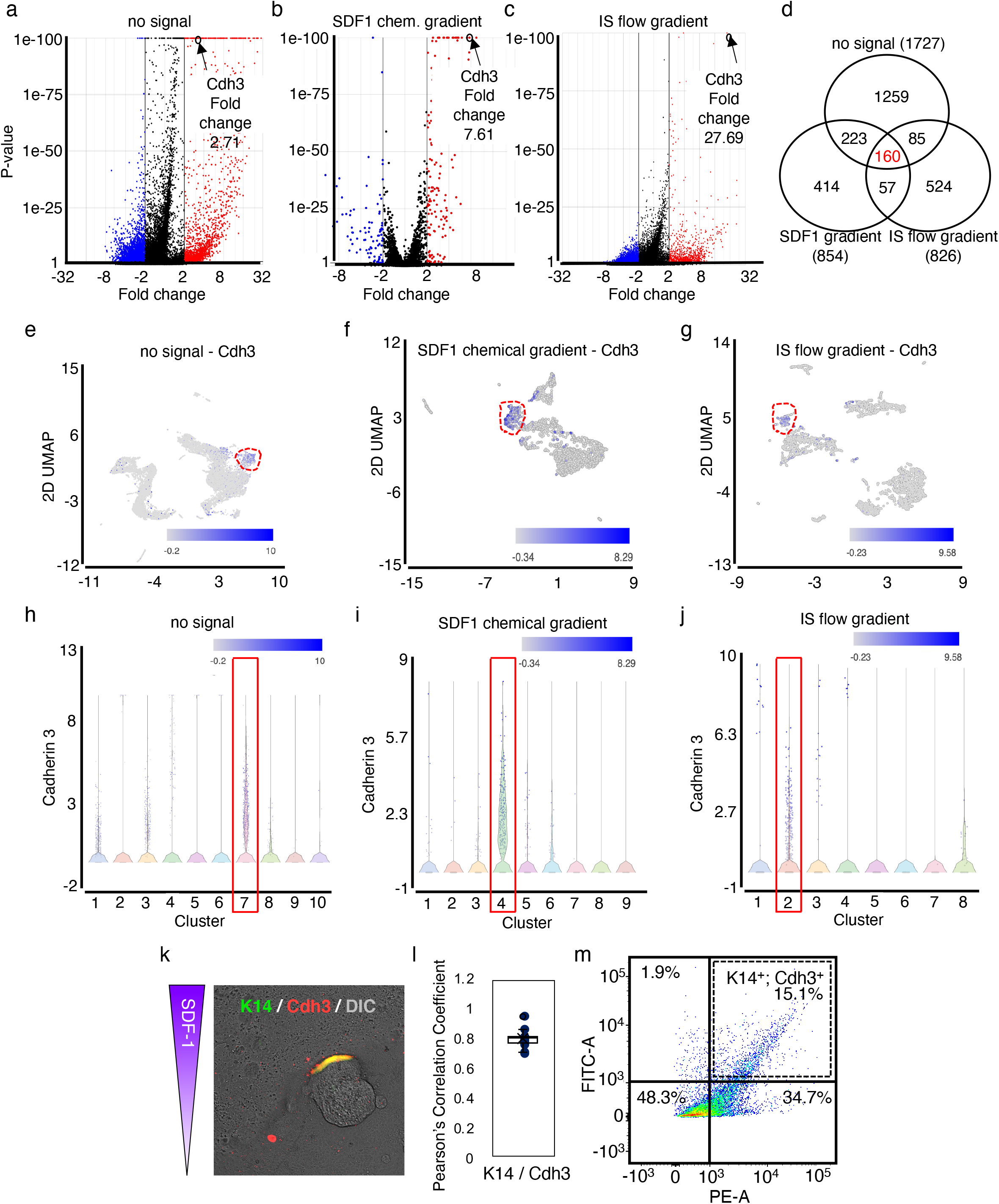
Cadherin 3 expression identifies a unique leader cell sub-population. **(a-c)** Volcano plots showing upregulated and downregulated genes for primary tumor organoids comparing the major K14+ population (#7 – no signal; #4 – SDF1 chemical gradient; #2 – IS flow gradient) to all other K14+ cell clusters within each respective sample: no directed migration (no signal), migration after SDF1 chemical gradient, and migration after interstitial (IS) flow gradient. Black arrows identify Cdh3 and its fold enrichment. **(d)** Genes found enriched in primary tumor organoids (no signal) were overlapped with genes enriched in primary tumor organoids following migration after SDF1 chemical gradient and migration after interstitial (IS) flow gradient in a Venn diagram. **(e-g)** UMAP plot and **(h-j)** violin plots showing Cadherin-3 (Cdh3) expressing cell cluster for no signal, SDF1 chemical gradient, and interstitial (IS) flow gradient. **(k)** Representative IF image of MMTV-PyMT tumor organoid after migration inresponse to SDF1 chemokine gradient showing Cdh3 (red) is present in K14+ (green) cells. **(l)** Pearson’s correlation coefficient analysis showing co-localization of K14 and Cdh3. **(m)** FACs sorting for K14+/Cdh3+ leader cells in MMTV-PyMT primary tumor organoids. For all experiments, *p<0.05 ANOVA with Tukey’s post hoc analysis.

To be able to test whether the K14^+^/Cdh3^+^ subpopulation of tumor cells could direct collective migration, we developed an assay to reconstitute invasive breast organoids and quantify directed collective migration potential (Fig. 4a). Organoids were isolated from normal breast of syngeneic age-matched non-tumor bearing mice. These are referred to as Mammary Gland (MG). When MGs were placed alone in the microfluidic device, in 3D collagen I and hypoxia, they remained viable and intact. In response to an SDF1 gradient there was no directed collective migration (Fig. 4b,c; quantified in 4d,e). Next, K14-Actin.GFP; MMTV-PyMT primary breast tumors were dissociated to single cells, Thy1^+^; Pdfgrβ^+^ CAFs were removed, and then K14^+^/Cdh3^+^ tumor cells isolated by FACS. This population of tumor cells was then mixed with normal breast organoids (MG) in ratios based on previously calculated percentages from primary tumor organoids (10% K14+ cells). Through immunofluorescent analysis, we confirmed in these reconstituted organoids that they contained a similar composition of K14+ cells as primary tumor organoids (Suppl. Fig. S3f). Added K14^+^/Cdh3^+^ cells were initially randomly distributed throughout the reconstituted organoids yet in response to an SDF1 gradient they polarized to the leading edge (Suppl. Fig. S3g), as was observed with WT PyMT breast tumor organoids(22). We refer to these reconstituted organoids as “MG + K14^+^/Cdh3^+^”.

**Figure 4.**
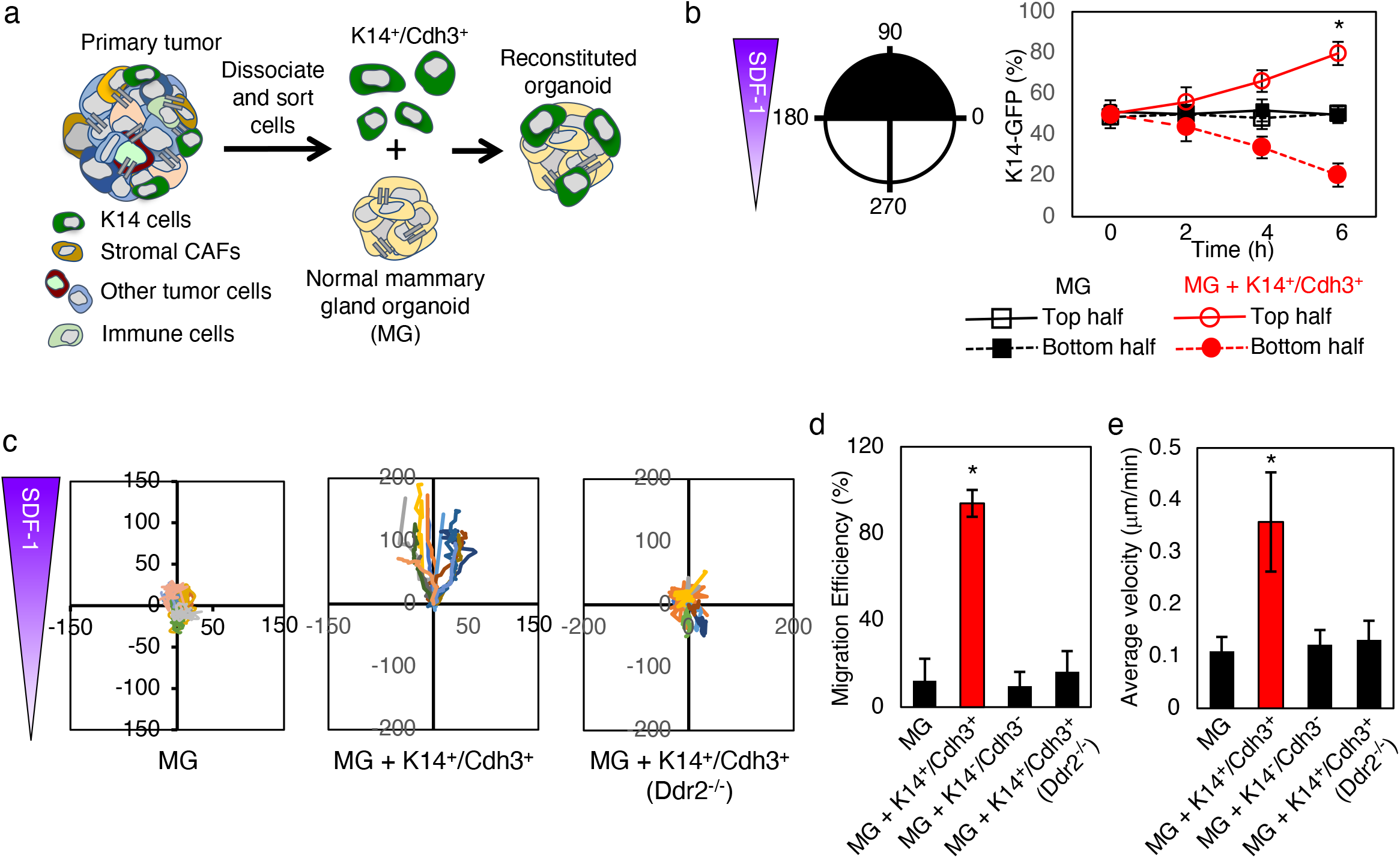
The K14+/Cdh3+ leader cell subpopulation can promote directed collective migration of non-migratory normal breast organoids to obtain. **(a)** Scheme of the organoid reconstitution assay: K14+/Cdh3+ leader cells are sorted from primary MMTV-PyMT tumors and mixed with normal mammary gland organoids. **(b)** Polarization of K14-Actin.GFP cells in reconstituted organoids after exposure to SDF1 chemokine gradient over a period of 6 hours. **(c)** Collective migration tracking maps for mammary gland organoids (non-migratory), K14+/Cdh3+ reconstituted organoids, and K14+/Cdh3+ (Ddr2^-/-^) reconstituted organoids. **(d-e)** Quantification of migration efficiency in the direction of the chemokine gradient and average velocity for reconstituted organoids. For all experiments, N=12-15 organoids within the microfluidic devices from a minimum of 3 different mice were analyzed, *p<0.05 ANOVA with Tukey’s post hoc analysis.

When “MG + K14^+^/Cdh3^+^” organoids were placed to the microfluidic platform (3D collagen I matrix and hypoxia), and exposed to an SDF1 gradient, directed collective migration now occurred in real-time (Fig. 4b,c; quantified in 4d,e). For collective migration of reconstituted organoids to occur required Cdh3+ tumor cells as when MG organoids were reconstituted with K14^low^/Cdh3^-^ tumor cells from sorted primary tumors, directional collective migration did not occur (Fig. 4d,e). The presence of the fibrillar collagen receptor discoidin domain receptor 2 (DDR2) in K14+ PyMT tumor cells has been shown to be required for directed collective migration in the MF device(22) and lung metastasis in vivo(36). Therefore, in other control experiments, MG organoids were reconstituted with K14^+^/Cdh3^+^ from Ddr2^-/-^; MMTV-PyMT primary breast tumors. In this experimental setting, directional collective migration did not occur (Fig. 4c; quantified in 3d,e). Deletion of Ddr2 in MMTV-PyMT tumors did not significantly affect the scRNAseq K14+ tumor cell profile nor Cdh3 expression in organoids (Suppl. Fig. S3h).

In sum, these results demonstrated that a K14^+^/Cdh3^+^ tumor cell subpopulation was necessary and sufficient to lead directed collective invasion of normal breast organoids through 3D collagen I matrices.

### Cadherin 3 expression is required for leader cell function

Cadherin 3 expression in cancer is associated with invasive behavior and poor clinical outcomes(37, 38) and Cdh3 overexpression in mesenchymal-like cell lines, that do not normally express Cdh3, can promote their collective cell migration(39). Immunofluorescent analyses (Fig. 3k,l) and scRNAseq analysis (Fig. 3e-j) of primary tumor organoids within our microfluidic system revealed that Cdh3 co-localized with K14+ cells at the leading edge and was not expressed by other “follower” tumor cells or associated stromal cells (CAFs, endothelial cell, myeloid cells). This unique enrichment of Cdh3 expression specifically in the putative tumor leader cell subpopulation led us to ask whether primary breast tumor leader cells require Cdh3 to lead collective invasion.

We identified multiple Cdh3 shRNA containing lentiviruses that efficiently (>90%) depleted Cdh3 in K14+ human (BT549) and mouse (4T1) invasive breast tumor cell lines (Suppl. Fig. S4a). Depletion of Cdh3 in both cell lines did not impact cell viability or proliferation (data not shown) but resulted in decreased invasion/migration of these cell lines through Matrigel (Suppl. Fig. S4b) and 3D collagen I gels (Suppl. Fig. S4c). These lentiviral vectors also co-express either GFP or RFP that allowed for identification, tracking, and quantification of transduced cells. We isolated breast tumor organoids from K14-Actin.GFP; MMTV-PyMT mouse breast tumors (Fig. 5), and human triple negative breast cancer (TNBC) patient xenografts (PDX – human in mouse breast tumors) (Suppl. Fig. S4) and then infected these with control (shSCR) or Cdh3 shRNA lentiviruses (mouse and human – two different shRNAs).

**Figure 5.**
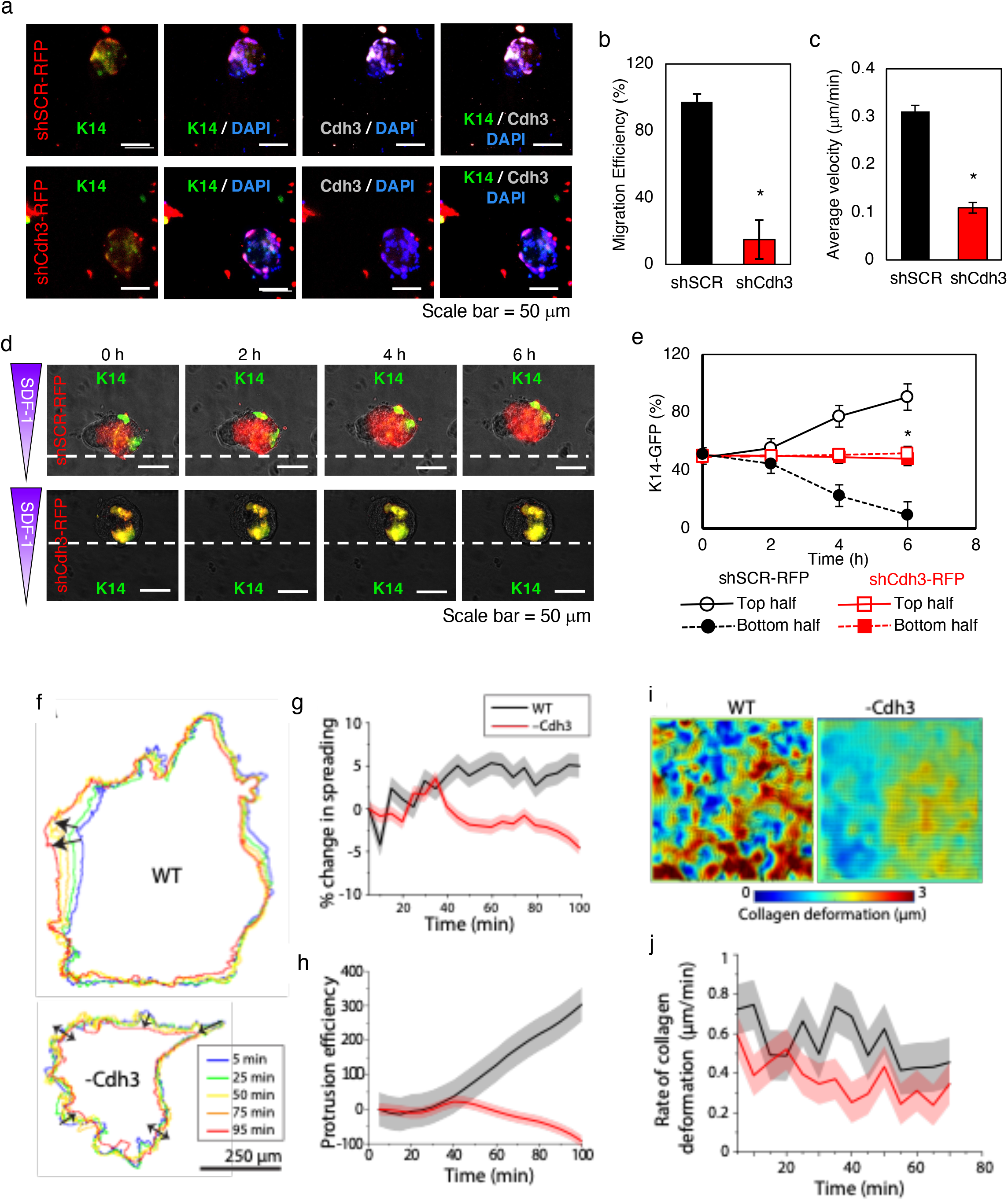
Cadherin 3 expression in leader cells is required for directed migration, K14+ cell polarization, protrusion efficiency, and collagen fiber deformation. (**a**) Representative IF images of shCdh3-RFP transduction (and corresponding scramble control) in K14+ (GFP) leader cells of primary tumor organoids. (**b-c)** Quantification of migration efficiency in the direction of the chemokine gradient and average velocity for shCdh3-RFP (and corresponding scramble controls). **(d)** Representative IF images of K14-GFP leader cell polarization (shCdh3-RFP and shSCR-RFP) after exposure to SDF1 gradient for 6 hours. **(e)** Quantification of K14+GFP polarization to the leading edge (shCDh3-RFP and scramble control) after exposure to SDF1 gradient for 6 hours. **(f)** Wild type (WT) and shCdh3 organoids encased in 2.3mg/ml collagen for 12h, live-imaging performed for 1.5h every 5 min interval, and protrusion boundaries traced with SiR-actin labeling. Protrusion outlines for representative organoids are plotted to demonstrate protrusion dynamics over time, visualize by the color-coded timestamps. Scale bar = 250 μm. Arrows denote unidirectional (single arrowheads) and regressive (double arrowheads) protrusions in WT and shCdh3 cases, respectively. **(g)** Percentage change in organoid area, relative to the reference timepoint, over time calculated from protrusion outlines. **(h)** Protrusion efficiency (%area-min) calculated in terms of area under the curve of percent spreading over time using Y=0 as baseline; here, regressive protrusions reduce protrusion efficiency. **(i)** Spatial heatmap of collagen deformation around the organoids obtained by subtracting positions of beads embedded within 3D collagen before and after trypsinization of organoids. **(j)** Rate of collagen deformation of live organoids (before trypsinization) over time. N=3 for WT and N=5 for shCdh3. In plots, solid lines represent mean and shaded background represents standard error (SE). For experiments in microfluidic devices (a-e), N=12-15 organoids from a minimum of 3 different mice were analyzed. For all experiments, *p<0.05 ANOVA with Tukey’s post hoc analysis.

Immediately after isolation, tumor organoids were placed in the microfluidic device (3D collagen I) under hypoxia and lentivirus delivered through fluidic lines (i.e., organoids are never exposed to tissue culture plastic). Successful lentiviral transduction and Cdh3 protein depletion in K14+ cells was confirmed through immunofluorescent analyses: K14 (Actin.GFP), lentiviral infection (RFP), and Cdh3 (Alexafluor-633) (Fig. 5a). Directed collective migration of transduced (RFP+) organoids was scored and analyzed. Depletion of Cdh3 resulted in decreased migratory efficiency and average migratory velocity (Fig. 5b,c). In Cdh3-depleted tumor organoids K14+ cells did not polarize to the leading edge (Fig. 5d,e). There was no change in organoid viability following depletion of Cdh3 and the number of K14+ cells per organoid did not change. Similar to mouse MMTV-PyMT breast tumor organoids, depletion of Cdh3 in human invasive TNBC PDX organoids also resulted in decreased migration efficiency and average velocity (Suppl. Fig. S4d-f).

In another approach, freshly lentiviral transduced MMTV-PyMT organoids (GFP+) were immediately placed in a standard breast tumor organoid culture: 3D collagen I gels containing a uniform concentration of either bFGF + EGF or SDF1 (i.e., no signaling gradient) and cultured under normoxic conditions(22). Under these conditions organoids send out circumferential invasive cellular projections that are led by K14+ cells but do not migrate collectively as a whole(19). We scored tumor organoids in which lentivirus derived GFP was expressed in the majority of cells for percentage of tumor organoid colonies with or without invasive phenotype (see representative examples (Suppl. Fig. S4g,h). Depletion of Cdh3 resulted in significantly decreased number of invasive tumor organoids (Suppl. Fig. S4i,j).

To determine if Cdh3 expression was required for in vivo lung metastases we performed syngeneic, orthotopic breast transplant experiments with mouse 4T1 breast tumor cells +/- Cdh3. 4T1 cells are “leader-like” in that they express both K14 and Cdh3. Although there was no difference in primary tumor growth, there were less lung metastases present in mice transplanted with Cdh3-depleted 4T1 cells (Suppl. Fig. S4k,l).

In summary, these results indicated that both mouse and human breast tumor leader cells required the presence of Cdh3 for polarization to the leading edge, their capacity to lead directed collective migration/invasion, and metastases in vivo.

### Leader cell polarization and function depend upon Cdh3-dependent protrusion stability and ECM collagen fiber deformation

Our computational model predicted that high protrusive forces in leader cells are required for net tumor organoid invasion. To experimentally test whether the presence of Cdh3 was important for protrusive activity and forces, we cultured control shSCR and Cdh3-depleted (shCdh3) mouse PyMT tumor organoids in 3D collagen I with embedded fluorescent beads and performed live imaging. We labeled F-actin and traced live protrusions along the periphery of the tumor organoids over time. Control WT tumor organoids stably protruded outward in a given direction (Fig. 5f). In contrast, Cdh3-depleted organoids were smaller in size and their periphery fluctuated back and forth, without stable directionality, over time (Fig. 5f). To quantify protrusion dynamics, we plotted percentage change in area of protrusions in tumor organoids over a ∼1.5 h tracking duration. While the protrusion area stably increased in WT tumor organoids, it fluctuated with an overall reduction in the Cdh3-depleted tumor organoids (Fig. 5g). We also calculated protrusion efficiency by integrating rate of change in protrusion area. This was higher for WT tumor organoid compared to the Cdh3-depleted ones (Fig. 5h). This experiment indicated that the presence of Cdh3 was important for stable protrusion formation and function in tumor organoids.

Next, we experimentally tested whether Cdh3 influenced tumor organoid interaction with the 3D collagen I ECM. We allowed tumor organoids to reside in 3D collagen I containing fluorescent embedded beads over 2 days and imaged bead displacement before and after trypsinization. We plotted net bead displacement and found that collagen deformations were dramatically reduced in Cdh3-depleted tumor organoids (Fig. 5i). We repeated these measurements for 4 different tumor organoids from multiple mice and averaged collagen deformation over time after trypsinization (Fig. 5j). This analysis further confirmed that Cdh3-depleted tumor organoids showed smaller collagen deformation indicative of lower forces. Similar changes in protrusion stability and collagen deformation were apparent when human BT549 cell spheroids (+/- Cdh3) were embedded in 3D collagen I (Suppl. Fig. S5a-d).

### Cadherin 3 regulates ECM feedback through Laminin 332 expression by leader cells

To determine how leader cell interaction with the ECM generates active ECM adhesion feedback, which is predicted to counteract ECM resistance at the leading edge, we noted that theCdh3+ subpopulation of K14+ leader cells were also enriched for hemidesmosomal receptor genes Col17 and Integrin α6β4. as well as their ligand Laminin 332, a basement membrane component (Fig. 6a,b). Immunostaining of invasive mouse MMTV-PyMT breast tumor tissue revealed that Laminin 332 was indeed present at the invasive leading edges, *in vivo* (Fig. 6c). Immunostaining of MMTV-PyMT tumor organoids for Laminin 332 and Collagen 17 demonstrated that Laminin 332 expression was observed in proximity to leader cells while Collagen 17 was present around the entire periphery of the organoids, possibly reflecting the fact that the extracellular domain of Collagen 17 is proteolytically released from cells(40) (Fig. 6d). After depletion of Cdh3 in MMTV-PyMT tumor organoids, expression of Laminin 332 was, surprisingly, significantly decreased compared to controls, whereas Collagen 17 expression did not change (Fig. 6d). Decreased Laminin 332 expression was also noted in Cdh3-depleted human PDX breast tumor organoids, without any change in Collagen 17 expression (Fig. 6e).

**Figure 6.**
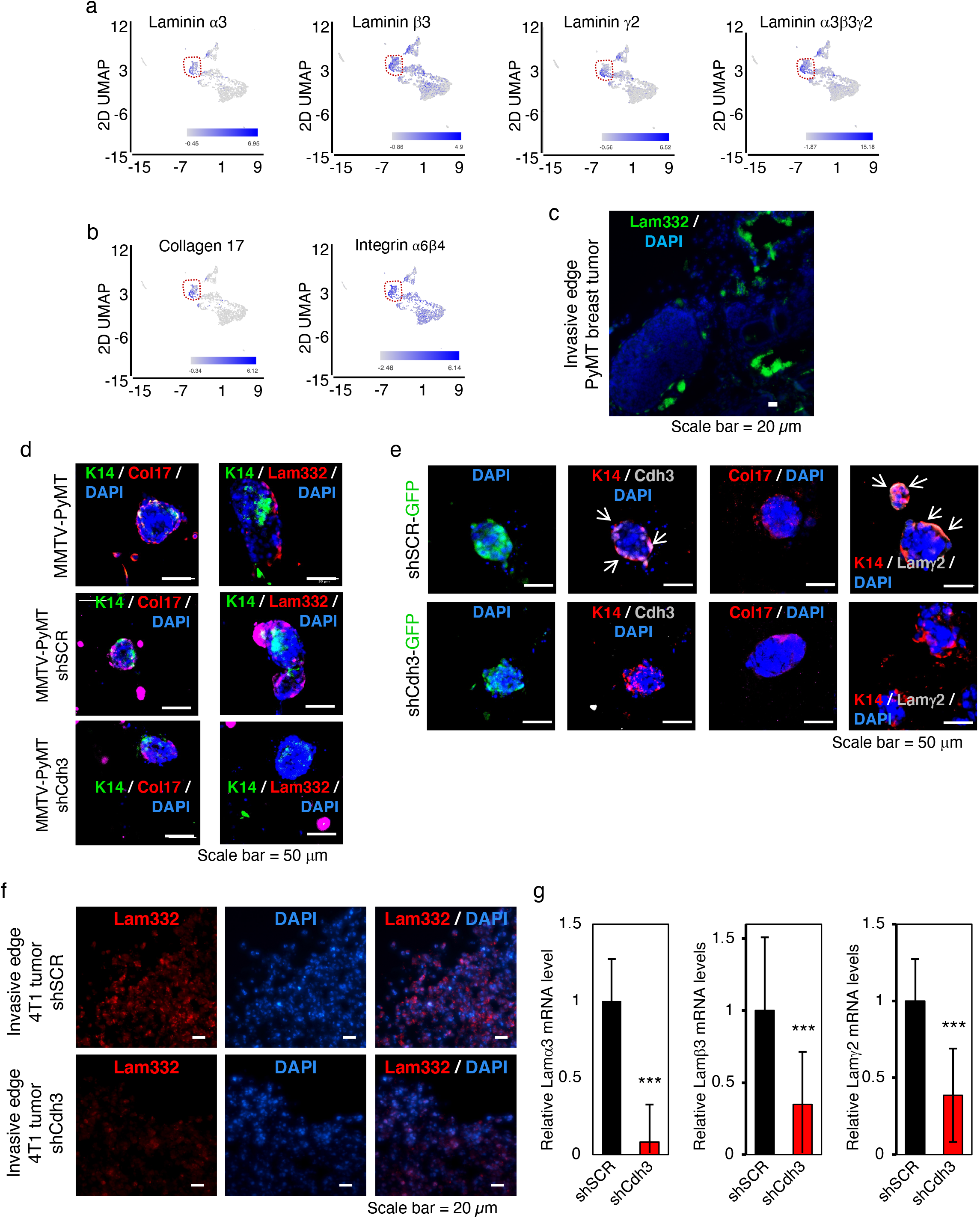
Cadherin 3 controls ECM feedback by regulating Laminin 332 expression in leader cells. **(a-b)** UMAP plots showing that the unique leader cell cluster #4, in an SDF1 chemical gradient, are enriched in Laminin 332 genes: Laminin α3, Laminin β3, Laminin γ2, and Laminin α3β3γ2 **(a)** and hemi-desmosomal receptors for Laminin 332 - Collagen 17α1 and Integrinα6β4 genes **(b). (c)** IF image of laminin332 at the leading edge in MMTV-PyMT primary breast tumor tissue section. **(d)** IF images of MMTV-PyMT tumor organoids (WT, shSCR-RFP, shCdh3-RFP) for K14-GFP, Collagen 17, and Laminin 332 expression. **(e)** IF images of human PDX tumor organoids (WT, shSCR, shCdh3) for K14-GFP, Collagen 17, and Laminin 332 expression. **(f)** IF images of the leading, invasive edge of control shSCR and shCdh3 4T1 breast primary tumor sections. **(g)** Expression of Laminin 332 genes (α3, β3, γ2) in control shSCR and shCdh3 4T1 breast primary tumors. For all experiments in microfluidic devices (d-e), N=12-15 organoids from a minimum of 3 different mice were analyzed. For all experiments, ***p<0.01 ANOVA with Tukey’s post hoc analysis.

In another approach, we used the syngeneic, orthotopic 4T1 transplant mouse model. Cdh3 depleted (or control: shSCR) 4T1 cells were implanted into the 4^th^ mammary fat pad of Balb/c females. Primary tumors were analyzed for Laminin 332 expression at the leading edge. Similar to the *ex vivo* primary tumor organoid experiment, Cdh3-depletion resulted in decreased expression of Laminin 332 at the leading edge (Fig. 6f), which was confirmed with mRNA analysis (Fig. 6g). These findings indicated that in primary tumors and tumor organoid leader cells the presence of Cdh3 controlled Laminin 332 expression suggesting that in leader cells at the leading edge of migrating tumors Laminin 332 interactions may contribute to computationally predicted ECM feedback.

### Cadherin 3 controls Laminin 332 transcription through β-catenin

To determine if Laminin 332 expression by leader cells was important for directed collective migration, we shRNA-depleted Laminin α3 in primary mouse tumor organoids (Suppl. Fig. S6a). Depletion of Laminin α3 resulted in decreased directional collective invasion (Fig. 7a-c), and K14 leader cells did not polarize to the leading edge (Fig. 7d). Immunostaining of human PDX organoids for Lamininγ2, mouse MMTV-PyMT organoids for Laminin 332, as well as the co-receptor Collagen 17 were performed (Fig. 7e). In human Laminin α3 depleted organoids and mouse Laminin α3 depleted organoids Lamininγ2 and Laminin 332 protein levels, respectively, were significantly decreased. Collagen 17 staining was not impacted as it was present around the entire periphery of the organoids (Fig. 7e).

**Figure 7.**
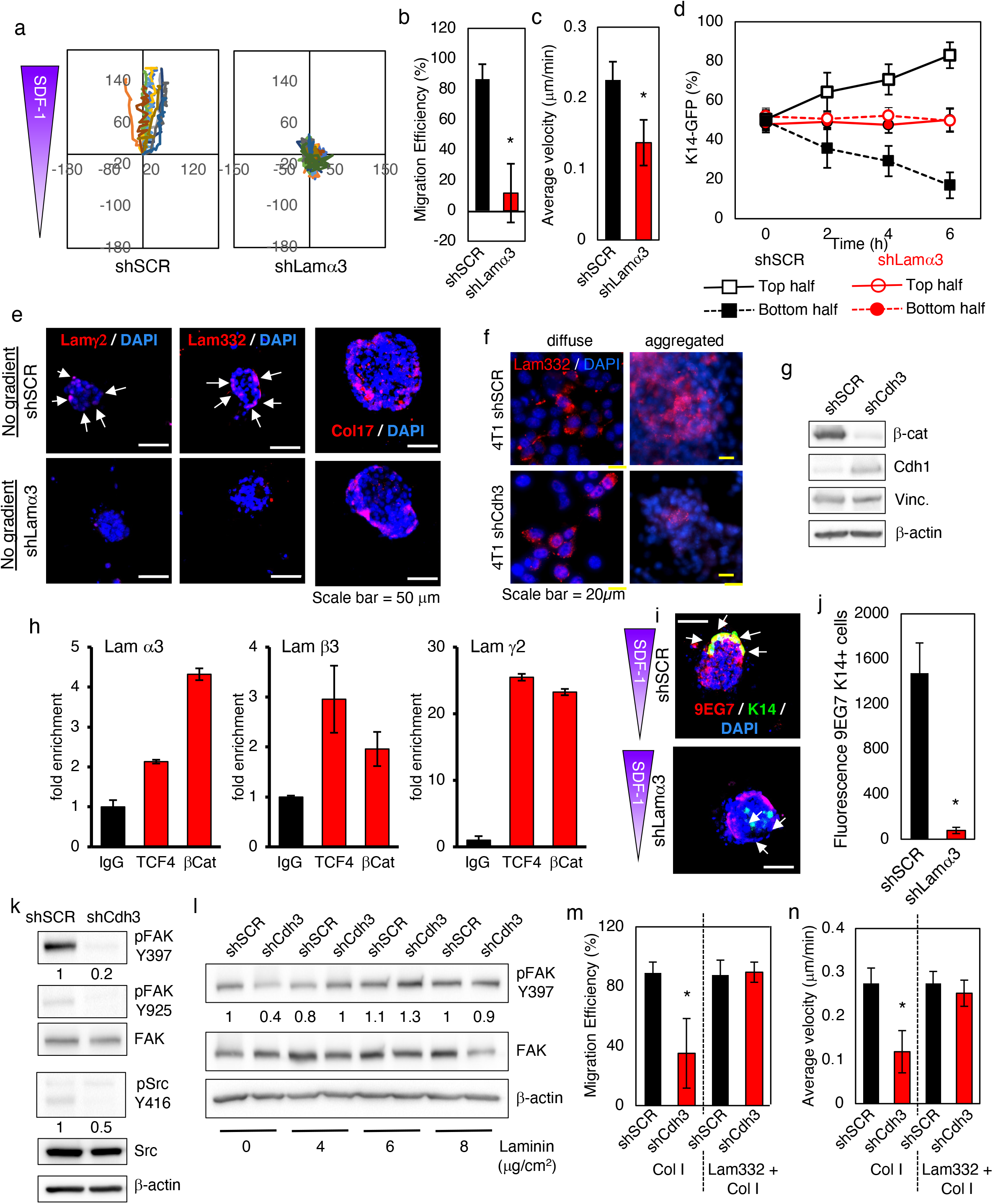
Cadherin 3 controls Laminin 332 transcription through β-catenin and Laminin 332 expression by leader cells regulates Integrin activity in leader cells. **(a-c)** Collective migration tracking maps, migration efficiency, and average velocity for shLamα3 and shSCR MMTV-PyMT tumor organoids after exposure to SDF1 gradients. **(d)** Quantification of K14+GFP localization (shLamα3 and shSCR) after exposure to SDF1 gradient for 6 hours. **(e)** Representative IF images of MMTV-PyMT tumor organoids (WT, shSCR, shLamα3) for confirmation of Laminin-α2, Laminin 332, and Collagen 17. Organoids have not been exposed to SDF1 chemokine gradient. White arrows delineate positive Laminin expression **(f)** IF images of 4T1 (shSCR, shCdh3) cells as diffuse or aggregated (cell contacted) cells for Laminin 332. **(g)** Western blot of confluent (cell contacted) 4T1 (shSCR, shCdh3) cells for β-catenin (β-cat), E-cadherin (Cdh1), and vinculin (Vinc.) **(h)** ChIP results for control IgG, anti-βCat, and anti-TCF4 antibodies on the promoter region of Lamα3, Lamβ3, Lamγ2 genes in confluent (cell-contacted) WT 4T1 cells. **(i)** Representative IF images of MMTV-PyMT tumor organoids (WT, shSCR, shLamα3) for K14-GFP, and activated integrin (9EG7) expression after exposure to SDF1 gradient. K14 leader cells are delineated by white arrows. **(j)** Quantification of total 9EG7 fluorescence in K14+ cells (shSCR, shLamα3). **(k)** Western blot of confluent 4T1 (shSCR, shCdh3) cells with the indicated antibodies. **(l)** Western blot of confluent 4T1 (shSCR, shCdh3) cells after adhesion to varying concentration of Laminin 332 ligand with the indicated antibodies. pFAK.Y397 quantification is indicated below the panel. **(m-n)** Migration efficiency in the direction of SDF1 gradient and average velocity of 4T1 spheroids (shSCR, shCdh3) encapsulated in Collagen 1 (2 mg/ml) and Collagen 1 (2 mg/ml) + Laminin 332 (0.1 mg/ml) matrices within microfluidic devices. For all experiments in microfluidic devices (a-e, i-j, m-n), N=12-15 organoids from a minimum of 3 different mice were analyzed. For all experiments, *p<0.05 ANOVA with Tukey’s post hoc analysis.

To determine the molecular mechanism whereby the presence of Cdh3 in leader cells regulated Laminin 332 expression we turned to the 4T1 mouse invasive breast tumor cell line as biochemical analyses in primary organoids is limited due to the small number of cells per organoid (200-500 cells per organoid). Laminin 332 was expressed by WT and Cdh3-depleted 4T1 cell when they were cultured diffusely (i.e., no cell-cell contacts) (Fig. 7f). To induce Cdh3-Cdh3 intercellular interactions, that would occur in a tumor in vivo or tumor organoid we grew 4T1 cells to confluence, where cells were in contact with one another (cell-cell adhesions). As opposed to control aggregated cells, aggregated 4T1 cells depleted of Cdh3 expressed significantly less Laminin 332 protein (Fig. 7f) and Laminin α3, β3, γ2 mRNA (Suppl. Fig. S6b). These results suggested that Cdh3-Cdh3 intercellular interactions were critical for the regulation of Laminin 332 transcription.

Cytoplasmic β-Catenin is recruited to the cytoplasmic tail of Cadherin receptor following cell-cell adhesion and as such can serve as a reservoir of stabilized β-catenin in contacted cells. β-Catenin also has the capacity to translocate into the nucleus where it can regulate gene transcription in a complex with the TCF family of DNA-binding proteins. Western blot analyses of confluent (cell-cell contacted) Cdh3-depleted 4T1 cells revealed that total cellular level of β-catenin protein was dramatically reduced (Fig. 7g). If the decrease in β-catenin level in confluent Cdh3-depleted 4T1 cells was due to its proteosommal degradation, as a result of the sequestering capacity of Cdh3 or lack thereof, we asked whether proteosommal inhibition could stabilize β-catenin cellular levels in 4T1 cells lacking Cdh3. This was indeed the case (Suppl. Fig. S6c). To determine if β-catenin/TCF nuclear complexes regulated Laminin 332 transcription we performed chromatin IP (ChIP) experiments to the Laminin 332 gene promoter regions in WT confluent 4T1 cells. As compared to control IgG ChIP, both TCF4 and β-catenin chromatin immunoprecipitation revealed the presence of these protein on the promoter region of all three Laminin 332 genes (Fig. 7h). Putative Laminin α3, β3, and γ2 promoter regions, predicted TCF binding sites, and DNA oligo primers used for ChIP analysis are shown in Suppl. Fig. S7. Positive control β-catenin and TCF4 ChIP to the SP5 gene promoter is shown in Suppl. Fig. S6d.

In sum these results indicated that in the presence of Cdh3-Cdh3 intercellular interactions β-catenin cellular protein levels are high, can translocate to the nucleus where it could regulate transcription of Laminin 332 genes. In the absence of Cdh3 there is increased β-catenin proteosommal degradation (decreased cellular β-catenin), and thus, less nuclear β-catenin to control Laminin 332 gene transcription.

### Laminin 332 expression by leader cells regulates Integrin activity in leader cells

Defective protrusion efficiency and traction forces in Cdh3-depleted, Laminin 332 low tumor organoids suggested that integrin-dependent focal adhesion function was altered. To determine if this is indeed true, MMTV-PyMT tumor organoids depleted of Laminin 332 were immune-stained with the 9EG7 antibody that recognizes activated forms of integrin β1. In Cdh3-depleted tumor organoids there was decreased 9EG7 staining in K14+ leader cells (Fig. 7i; quantified in 7j). There are no changes in the total cellular levels of b1 Integrin in primary organoids in the MF device(41). In confluent 4T1 cells depleted of Cdh3, FAK and Src activation were significantly reduced (Fig. 7k). To test whether this could be due to the lack of Laminin 332 we added Cdh3-depleted aggregated 4T1 cells to Laminin 332 coated plates. FAK activation was rescued (Fig. 7l). In another approach we placed aggregated 4T1 spheroids (SCR and Cdh3-depleted) in our microfluidic platform within Collagen I or Collagen I + Laminin 332 (1:1) matrices and scored migration efficiency and velocity. As anticipated Cdh3-depleted 4T1 spheroids did not collectively migrate in Collagen I (Fig. 7m,n; Suppl. Fig. S6e). However, when placed in Collagen 1 + Laminin 332 containing device Cdh3-depeleted 4T1 spheroids now efficiently collectively migrated in a directional manner (Fig. 7m,n; Suppl. Fig. S6e). Images of embedded 4T1 spheroids within the devices revealed that 4T1 aggregates depleted of Cdh3 were less densely packed and occasional single cells exited, reflecting weakened cell-cell interactions when Cdh3 was depleted (Suppl. Fig. S6f).

In sum these results indicated that Cdh3 regulates Integrin/FA activity in leader cell protrusions by controlling the local production of Laminin 332.

## Discussion

Although there has been significant progress towards understanding molecular and functional heterogeneity of tumor cells(42, 43) and associated stromal cells(35) within the tumor microenvironment, leader cell heterogeneity, specifically, has not been addressed particularly in primary tumors. Using primary breast tumor organoids from a highly invasive, metastatic mouse GEMM and an invasive, metastatic TNBC human tumor PDX model, we observed multiple populations of histologically-defined leader cells, that varied depending upon the environmental signal. From analysis of scRNAseq data we were able to identify and isolate a subpopulation of K14+ leader cells, defined by the enriched expression of Cadherin 3. The presence of Cdh3 in this subpopulation of leader cells was critical for collective migration

This subpopulation of K14+ leader cells identified in response to an SDF1 chemical gradient is also enriched for a number of other cell surface signaling receptors, such as Fgfr2, Jag1, Jag2, that have been associated with breast cancer progression and metastases(44),(45, 46),(47). Whether they also control leader cell function(s), and if so, how remains to be determined. The Epithelial Mesenchymal Transition (EMT), or EMT inducing transcription factors per se, have also been implicated as controlling breast tumor invasion and metastasis. Of the major tumor EMT transcriptional regulators (e.g., Snail, Twist, Zeb, and Prrx families) only Snail2 was enriched in this leader cell subpopulation. Expression of others were predominantly enriched in CAF subpopulations. Snail2 enrichment is particularly interesting as it has recently been shown to transcriptionally activate expression of Cdh3 and requires Cdh3 for its EMT actions(48).

Cadherin-mediated cell-cell interactions between follower cells, between leader cell, between leader and follower cells, as well as between CAFs and leader cells are a central component of collective cell migration(49, 50). In tumor cells the dynamic expression of E-cadherin, or Cdh1, has received most attention but epithelial tumor cells can switch from expressing E-cadherin (Cdh1) to P-cadherin (Cdh3)(51, 52). High Cdh3 expression in patients with a variety of cancers, including invasive TNBC breast cancers, correlate with poor prognosis(37, 51, 53, 54). Cdh3 is also expressed by normal basal breast epithelial cells(52). Developmental genetic defects are associated with loss of Cdh3 and these have been linked to altered cell-cell interactions and cell-matrix adhesions(52). Here we find Cdh3 expression to be selectively enriched in bone fide leader cells of primary breast tumors. Within this leader cell subpopulation Cdh1 is also expressed but at lower levels than other tumor cells. In a mouse C2C12 myoblasts cell line (which does not endogenously express Cdh3) exogenous Cdh3 expression, but not Cdh1 expression, conferred the capacity for them to undergo collective migration(39). Some models of collective migration suggest leader cell-follower cell boundaries consist of heterotypic Cdh1-Cdh3 junctions(49). Whether a Cdh1-Cdh3 heterotypic interaction plays a role in primary tumor organoid collective migration, and if so, how remains to be determined.

We find that that the presence of Cdh3 in leader cells is required for the efficient protrusive activity, and as such, the ability for leader cells to polarize to the leading edge. The observed preferential polarization of Cdh3 leader cells towards the tumor organoid front suggests that leader cells generate higher protrusive forces compared to the follower cells. We started our model simulations with this premise of differential forces and polarity, however, higher forces in leader cells as the only mechanism required for their observer frontward polarization is not sufficient. With weak leader-follower adhesions, leader cells escaped the organoid into the surrounding collagen. Conversely, if leader cell adhesions with followers are made stronger, they stay adhered to followers and are unable to engage with the ECM which reduces their ability to invade the surrounding collagen.

As leader cells come to the leading edge they engage with the ECM (mostly fibrillar collagens) and subsequently cause the whole tumor organoid to move collectively. This suggests that leader cells actively engage with collagen and reduce the resistance presented by the collagen through an active adhesion feedback. This ECM feedback is triggered only upon leader cell engagement with the ECM, likely cause a rise in leader cell protrusive forces that would be transmitted to follower cells according to force equilibrium. When follower cells engage with the ECM, this feedback does not appear to arise. Thus, in our model, active ECM feedback benefits the Cdh3+ leader cells directly and could increase follower cell forces indirectly via force transmission. This active feedback could arise from alignment of collagen, degradation of collagen, or increased forces in leader cells due to integrin-based signaling, or a combination thereof. Our computational model used energy-based criterion to implement this active feedback, which does not explicitly distinguish among these possibilities. We did however observe experimentally that Cdh3-Cdh3 interactions in leader cells at the leading edge controlled local production of Laminin 332 by these cells and that Laminin 332 production or presence was required for efficient Integrin/FA mediated protrusion and force transduction.

Cadherin-based cell-cell adhesion have been shown to impact cell-ECM adhesions and traction force (55) (56), but how cadherin-based mechanical coupling between cells leads to increased ECM force transmission is not fully understood. Overexpression of Cdh3 in a myoblast cell line induces expression and secretion of decorin which modifies collagen fiber organization and alignment, and thus (57). This altered collagen structure results in Integrin activation, and decorin was required for Cdh3-induced collective migration. Thus, in this setting Cdh3 indirectly stimulates ECM traction forces. Herein, we find that Cdh3 positive primary tumor leader cells do not express decorin. Rather, decorin was expressed by organoid associated CAFs, which are in close proximity to the leader cells(22). Whether CAF produced decorin is required for Cdh3 mediated ECM traction force generation by collectively migrating tumor organoids remains to be determined. Regardless, herein, in primary tumor cells as opposed to cell lines, we find that Cdh3+ leader cells in tumor organoids indirectly stimulate ECM traction forces through the production of the hemidesmosome ligand Laminin 332. Cdh3 has also been shown to interact with β4 Integrin, presumably in cis, and affect cell surface α6/β4 Integrin levels(58). Since α6/β4 Integrin is one of the cell surface receptors for Laminin 332 this may also contribute to Cdh3 action in leader cells.

Perhaps hemidesmosome function in leader cells also contributes to the traction forces generated by tumor organoids. We found that the presence of Cdh3 in leader cells controls the transcription of Laminin 332. Laminin 332 is a major basement membrane ligand for hemidesmosomes and we show that its expression is enriched in the Cdh3+ leader cell subpopulation in vivo and in tumor organoids invading in 3D collagen I gels. Hemidesmosomes are cell matrix adhesive structures that associate with the keratin cytoskeleton, particularly in pseudo-stratified epithelia such as skin. In skin they anchor the basal layer of epithelial cells to the basement membrane through an association with Laminin 332(59). The hemidesmosome intermediate filament association enables cells to withstand mechanical stress and tension(60) and recently were shown to modulate cellular mechanical force through intracellular crosstalk with FAs(61). In keratinocytes loss of hemidesmosomes resulted in increased traction forces generation and cell spreading. But herein in primary breast tumor organoids loss of Cdh3, and as a result, decreased Laminin 332 expression resulted in decreased traction forces in 3D collagen I matrices. Hemidesmosome function can be cell and context dependent, as deletion of plectin (a cytosolic adapter associated with hemidesmosomes) in myoblasts and fibroblasts results in decreased traction forces(62, 63). Although the subpopulation of leader cells we identified are enriched for the expression of hemidesmosome components, as well as other genes typical of basal keratinocytes and basal breast myoepithelial cells, in the context of an invasive tumor they perhaps behave in a more mesenchymal manner as evident by the enriched expression of Snail2 and Cdh3.

## Materials and Methods

### Computational simulation of leader cell polarization and function in directional collective migration

We used a Cellular Potts approach to model heterogeneous tumor organoid migration in 3D because it is efficient in defining deformable 3D cell bodies. We model a collection of biological cells by attaching to each lattice point (pixel) (*i*; *j*) of a square lattice a label *σ*_*ij*_, which identifies the corresponding cell, and a label *τ*(*σ*_*ij*_), which identifies cell type. Adjacent lattice sites are defined to lie within the same cell if they have the same value of *σ*_*ij*_. The system evolves by the random movement of individual pixels subject to transition probabilities based on the Monte Carlo method(64). At each Monte Carlo Step, two neighboring pixels are chosen randomly, with one as source pixel and the other as target pixel. If both pixels belong to the same cell (*i. e*., *σ*(*source*) = *σ*(*target*)), then no changes are made to the lattice. Otherwise, the source pixel attempts to occupy the target pixel based on Monte Carlo acceptance probability, which is calculated from the difference in total system energy. We evaluate the total system energy associated with the configuration before the move and after the move as per the following equation:

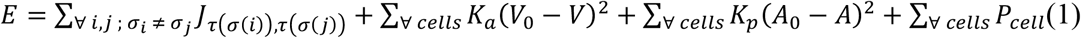

Here, *J*_*τ* (*σ*(*i*)), *τ* (*σ*(*j*))_ represents contact energy for the two cell types in contact *τ* (*σ*(*i*)), *τ* (*σ*(*j*)). The first term represents contribution from the total energy due to cell-cell adhesions. Second and third terms represent contributions from bulk elasticity of the cell and cell-surface contractility, respectively. *K*_*a*_ and *K*_*p*_ are constants for bulk elasticity and contractility, respectively. *V*_0_ and *A*_0_ are target volume and surface area that the cell has in isolation. *P*_*cell*_ represents effective protrusive energy of a cell (discussed ahead). After calculating energy of system before (*E*_*i*_) and after (*E*_*f*_) the copy attempt will always be successful if *E*_*f*_ < *E*_*i*_, *i. e*. Δ*E* < 0. If Δ*E* ≥ 0, the copy attempt is accepted with a probability of 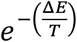. Higher values of *T* would tend to accept more unfavorable copy attempt. Here, we extend the Potts framework to model cellular invasion. We assign an effective protrusive energy to each cell, dependent on its cell type and interaction with its neighboring cells and ECM (spaces not occupied by the cell are assumed to belong to ECM).

Organoids were modeled as a spherical collection of cells with 10% leader cells (green) mixed randomly with non-leader “follower” cells (blue) (Fig. 1a,b). Adhesions are annotated as dotted lines, where *J*_*FF*_ represents follower-follower adhesion energy (cyan) and *J*_*LF*_ represents leader-follower adhesion energy (red). Based on our experiments done on primary organoids in a uniform chemokine gradient, we assume that leader cells sense the defined chemokine gradient and extend protrusions in that direction. We define leader cell protrusive force *P*_*L*_ (a dimensionless quantity) as following:

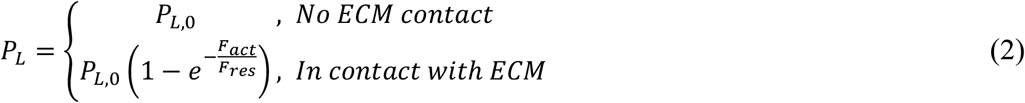

Here, *P*_*L*,0_ is the inherent protrusive force of leader cells without any feedback from the ECM. When leader cells encounter the ECM, such as an organoid invading into collagen, there are two counteracting mechanisms that alter this effective protrusive force produced by leader cells. First, the ECM serves as a steric barrier causing a resistance force *F*_*res*_, which lowers effective leader cell protrusive force due to ECM density and crosslinking. Second, cell-ECM adhesions are expected to activate mechanotransductive signaling and remodel the collagen matrix via degradation or fiber alignment. These leader cell-ECM interactions are assumed to produce an active force *F*_*act*_ that works against the ECM resistance force. Without this active force, ECM resistance would simply stop any invasion due to passive ECM resistance. We write this active force as:

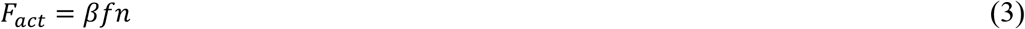

Here, *n* is the fraction of leader cells engaged with the ECM (*β∼*1) and *f* is force per leader cell. 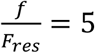. The degree of ECM feedback *β* is defined as a sigmoidal Hill function in term of the area of the leader cell in contact with the ECM as following:

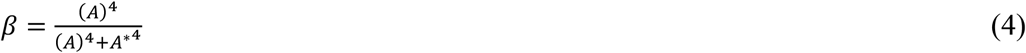

Here, *A* is the area of the leader cell in contact with the ECM relative to the total cell area and *A*^*∗*^ = 0.05 is a calibration constant. Based on this function, *β* = 0 when leader cells are within the organoids and *β* quickly rises to 1 as it interacts with the ECM, thus causing the active ECM feedback defined in Eq. 3.

In our model, follower cells are assumed not to follow the chemokine gradient, i.e., they are permitted to randomly polarize in any direction. We also assume an overall mechanical and chemical communication between follower and leader cells. Thus, follower cell protrusive force depends on average leader cell force, defined as:

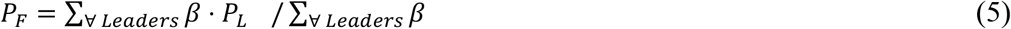

This average *P*_*F*_ is assigned to all follower cells.

### Primary breast tumor organoid isolation and culture

MMTV-PyMT mice (FVB/n) were obtained from The Jackson Laboratory and crossed to K14-Actin.GFP mice (FVB/n) (a transgenic mouse in which an Actin.GFP fusion protein is expressed under the control of the keratin-14 promoter), which is functional in mitotically active epidermal cells(65), to generate K14-GFP tagged MMTV-PyMT mice. They express Actin.EGFP only in K14+ cells. The endogenous K14 gene is not altered in these mice. We refer to all K14-positive cells obtained from this mouse as “K14-GFP.” Tumor-bearing mice were monitored weekly and euthanized at 12 weeks. All mice were used in compliance with the Washington University’s Institutional Animal Care and Use Committee and approved under protocol #20150145.

Mouse primary mammary tumor organoids were obtained as previously described(8), immediately mixed with collagen I (2 mg/ml, Corning) solution and loaded into the middle chamber of a previously described and characterized microfluidic device(22). Organoids are roughly 200-500 cells each. Collagen was allowed to polymerize (37°C, 1 hr, 20% O_2_), and media (DMEM, 10% FBS, P/S) delivered to the top and bottom fluidic lines and cultured in 2% O_2_ for 48 hours. At no time were the isolated tumor organoids exposed to plastic tissue culture dishes.

In other experiments we used “standard breast tumor organoid cultures”(8). Briefly, tumor organoids were embedded in 2 mg/ml collagen 1 solution (2-3 organoids/µl), and 100 µl suspension was plated in 4-well coverslip-bottomed chambers. Samples were placed on a 37°C heat block (30 min), followed by incubation at 37°C (30 min) to allow polymerization, and cultured in 1 ml of 2.5nM FGF2 or SDF1 in organoid media under normoxic conditions(18).

Patient derived TNBC metastatic xenograft (PDX) donor mouse models were obtained from JAX laboratories (JAX J000080739) and serially passaged through breast implantation into the fourth mammary fat pads of immunodeficient gamma (NSG) mice. Primary breast tumor tissue was minced, resuspended in Matrigel and injected mice. Tumor growth was monitored weekly and visible after eight weeks. When tumors reached @ 1 cm, mice were euthanized and organoids obtained. These organoids were then handled and treated in the same manner as the mice mammary tumor organoids, described above.

For studies using Laminin 332, we encapsulated 4T1 spheroids in a 1:1 ratio of collagen I (2mg/ml) to Laminin 332 (0.1 mg/ml, Biolumina). To promote directed collective migration, we induced a chemokine chemical gradient of SDF1 (0.05 ug/ml, Sigma-Aldrich) or interstitial fluid flow gradient (12 µm/s) via the top fluidic line(22).

### Single cell RNA sequencing (scRNAseq) and analysis of breast tumor organoids

Primary breast tumor organoids were isolated from the microfluidic devices after 48 h or hypoxia (no signal) or after promoting collective migration within microfluidic devices in response to a SDF1 chemokine chemical gradient or interstitial (IF) flow gradient; 18 hours; average 150 µm length of directional migration(22)). Organoids were extracted from the device and isolated down to the single cell level by delivering a low concentration collagenase/trypsin digestion. After successful extraction, cells were stained for live/dead (FxCycle Violet; ThermoFisher), and live cells sorted via FACs (BD FACSAria II). After sorting, cells were centrifuged and resuspended in scRNA sequencing buffer and final concentration was adjusted to 1000 cells/ul and sent for scRNA processing (10X Genomics). Samples were processed with each library sequenced on 0.125 NovaSeq S4 flow cell (300 cycles; 1.000 flow cell total). Read alignment, gene expression estimation, normalization and quality control were performed using the Cell Ranger Single-Cell Software Suite (10x Genomics). Cell Ranger count was used to align samples to the reference genome (mm10), quantify reads and filter reads with a quality score below 30.

All single cell RNA-sequencing (scRNAseq) data were analyzed using Partek Flow software. For quality control, nuclei with mitochondrial content >5% were removed. Nuclei that are doublets or multiplets were filtered out by two steps. First, nuclei with more than one marker gene expressed were removed. Then cells with high UMI and gene number per cell were filtered out. UMI counts were normalized following Partek Flow recommendations: for each UMI in each sample the number of raw reads was divided by the number of total mapped reads in that sample and multiplied by 1,000,000, obtaining a count per million value (CPM), the normalized expression value was log-transformed (pseudocount = 1). Starting from the normalized data node, we performed clustering analysis for each sample separately by means of graph-based clustering which employs the Louvain algorithm. Clustering analysis was done based on the first 100 principal components. To visualize single cells in a two-dimensional space, we performed a uniform manifold approximation and projection (UMAP) dimensional reduction using the first 20 principal components for each sample separately and for the entire data set.

We identified marker genes for each identified cell group by computing ANOVA comparing each cluster to all the other cells in the data set to obtain a list of genes with fold change (FC) >2 ranked according to ascending p-value. Using the list of genes generated for each cluster, we compared with known gene markers for various cell sub-types based on recently published data sets(19, 35, 66, 67). We also performed differential expression of genes between cell clusters within a sample using ANOVA analysis and considered genes with FC>2 or FC<-2 and p-value <0.05 as differentially expressed.

### Breast tumor cell line culture

BT549 and 4T1 cells were purchased from the American Type Culture Collection (ATCC; Masassas VA) and cultured in Dulbecco’s modified Eagle’s medium (DMEM) with 10% serum and penicillin-streptomycin. Cells were routinely checked for the presence of mycoplasma by polymerase chain reaction (PCR) amplification using primers Myco + (5′-GGG AGC AAA CAG GAT TAG ATA CCC T-3′) and Myco− (5′-TGC ACC ATC TGT CAC TCT GTT AAC CTC-3′) every 6 months. Only mycoplasma-negative cells were used in these studies. 4T1 cells were cultured either diffusely – non-contacted (30% plate coverage) or confluent – cell-cell contacted. To make 4T1 cell spheroids (aggregates) for 3D culture, cells were cultured on low adhesion plates (Corning) to naturally form spheroids, single cells were filtered out, and then spheroids loaded into the microfluidic devices as with MMTV-PyMT breast tumor organoids.

### shRNA lentiviral depletion of gene expression

We synthesized 5 different shRNA lentivirus particles per gene - both human and mouse specific Cdh3 (Origene, TR30023 (human), TL500332 (mouse)) and Laminin α3 (Sigma-Aldrich), as well as corresponding scramble (shSCR) controls. shRNA sequences used are listed in Supplemental Table 2. Each lentivirus also expressed either GFP or RFP that allowed for identification and quantification of transduced cells. Human BT549 breast tumor cells and mouse 4T1 breast tumor cells were exposed to transduction media, treated with selection media (puromycin), and successful transduction confirmed by visualized GFP expression. BT549 and 4T1 cells began to express GFP after 48 hours; at that time point, cells were lysed and Western blot was performed to confirm successful knockdown. In each case, multiple shRNAs were identified that decreased protein expression >80%.

Mouse MMTV-PyMT primary breast tumor organoids and human PDX primary breast tumor organoids embedded in 3D collagen I within the microfluidic devices were transduced with shRNA expressing lentiviruses directly via fluidic lines. Organoids were exposed to transduction media for 16 hours, and successful transduction confirmed by visualizing fluorescence marker expression. Organoids began to express GFP/RFP after 18 hours. Organoids were stained for Cdh3 and Laminin 332 expression to confirm successful knockdown.

### Transwell Invasion Assay

Transwell invasion assays were performed using a 24-well polycarbonate membrane (Corning) with 8.0 µm pore size. The upper chambers were precoated with 1 mg/ml Matrigel and incubated at 37 °C for 2 h. The cells (5 × 10^4^/well) in DMEM containing 1% FBS were seeded into the upper chambers, and 600 ul of DMEM with 10% FBS was added to the lower chambers. After incubation for 24 h at 37 °C, polycarbonate membranes were stained with HEMA3 staining kit (Fisher). BT549 or 4T1 cells on the upper surface were removed with cotton-tipped swabs. The number of invasive cells were counted from six randomly selected visual fields using compound light microscopy (200x magnification).

### 3D Cell Invasion Assay

For 3D cell invasion assays, 10^5^ cells (BT549 or 4T1) were embedded in 20 μl of type I collagen gel (2.2 mg/ml, Corning). After gelling, the plug was embedded in a cell-free collagen gel (2.2 mg/ml) within a 24-well plate. After allowing the surrounding collagen matrix to gel (1 h at 37°C), 0.5 ml of culture medium was added to the top of the gel and cultured for another two days. Invasion distance from the inner collagen plug into the outer collagen gel was quantified.

### 4T1 syngeneic orthotopic breast transplants

Eight-week old female BALB/cJ mice (Jackson Labs) were anaesthetized with a ketamine/xylazine cocktail (90 mg/kg, 1 ketamine and 13 mg/kg, xylazine, intraperitoneal injection) and the abdomen was sterilized using povidone iodine (Betadine) solution and ethanol. A small Y-shaped incision was made in the lower abdominal skin to expose the fourth mammary gland using surgical scissors and bleeding vessels were cauterized. 4T1 cells (1 × 10^6^) in 50 µl DMEM were injected into the fourth mammary gland using a 29-gauge needle. The skin flaps were replaced and closed using 9mm wound clips, and the surgical site was swabbed with triple-antibiotic cream. Primary tumors and lungs were collected at end stage (primary tumor @ 2cm) and processed for immunohistochemistry. The lungs and primary tumors were then fixed in 10% formalin for 24 h, cryopreserved in 30% sucrose overnight, and finally embedded in OCT and frozen in a dry ice/ethanol bath. Lungs were processed in three step-sections of 50 um with two serial sections of each step. Metastatic tumor nodules were counted and averaged per lobe of lung (5 lobes) per mouse. Primary breast tumors were stained for Laminin 332, and gene expression for Cdh3 and Lamα3, Lamβ3, and Lamγ2 determined via QPCR.

### Live-cell imaging and analysis

After culturing organoids for 48 hours in 2% O_2_, we induced collective migration and performed live-cell imaging (Nikon Ti-E, 10x, 40x, 60x; controlled temperature, humidity, and oxygen (2% O2)). Each organoid within the device was marked using Nikon Imaging software, and pictures were taken every 20 minutes for a maximum of 18 hours. After imaging, devices were used for immunofluorescence labeling and imaging, or organoids were extracted from the device for single cell sequencing analysis.

Image analysis was performed using Metamorph, Matlab, and FIJI to quantify organoid migration efficiency (%) in the direction of the gradient(22), average velocity (µm/min), and direction of travel (migration maps). We also tracked and quantified K14-GFP localization over time. At various time points, images of organoids were divided into top (front; direction of migration) and bottom (back) halves, and total K14-GFP fluorescence of each half was calculated using FIJI and the following formula: cell fluorescence = integrated density – (area of half x mean fluorescence of background).

### Immunofluorescence

All immunostaining was performed after imaging studies with organoids maintained within the devices, and all reagents were delivered via microfluidic lines. After fixing and blocking, organoids were stained for Cadherin 3 (ThermoFisher), Collagen 17 (ThermoFisher), activated Integrin β1 (9EG7; BD Biosciences), mouse Laminin 332 (gift from Dr. Takako Sasaki, OITA University), or human Laminin γ2 (EMD Millipore); all primary antibody staining was incubated overnight at 4°C. Species-specific secondary antibodies (488, 555, or 633 wavelength) and nuclei staining (DAPI) were also used. Imaging was performed via confocal microscopy (Zeiss, 63X). All antibodies used are listed in Supplemental Tables 3 and 4.

Tumor tissues were fixed with 10% formalin for 24 h, embedded in paraffin and sectioned (5 μm). After dewaxing using a graded alcohol series and antigen retrieval, the tumor slides were treated with 3% hydrogen peroxide, and then blocked with 10% normal goat serum for 1 h. The slides were incubated with the primary antibodies against Laminin-α3 (1:500) at 4°C overnight. After three washes with PBS, the tissue sections were incubated with goat anti-rabbit AlexaFluor 594 antibody (Invitrogen) for 2h at 4°C. Slides were then mounted using VECTASHIELD (Vector Laboratories) with DAPI and coverslips applied.

### Gene expression

Total RNA was extracted from cells or primary tumor tissues by RNeasy Plus Mini Kit (Qiagen). Subsequently, RNA was reverse transcribed into cDNA using the SuperScript First-Strand Synthesis System for RT-PCR (Invitrogen). cDNA was amplified using SYBR Green PCR Master Mix (Applied Biosystems) on QuantStudio 3 Real-Time PCR System (Applied Biosystems). The primers targeted Lamα3, Lamβ3, Lamγ2, Cdh3 and β-actin are listed in Supplemental Table 5. The thermocycling conditions were as follows: initial denaturation at 95°C for 10 min, 40 cycles of 95°C for 15s, 60°C for 60s. β-actin was chosen as an endogenous control. The 2^−ΔΔCT^ method was used to calculate the relative mRNA levels.

### Protein expression

Cells were lysed in RIPA buffer plus protease inhibitors (Sigma-Aldrich). Protein concentration was measured using Bradford Reagent (Biorad). Lysates were subjected to SDS-PAGE, transferred to PVDF membranes, blocked in 5% milk, incubated with primary antibody overnight, secondary antibody for 2 hours and visualized using SuperSignal WestPico and/or Super Signal West Femto Chemiluminescent Substrates (ThermoFisher Scientific). Exposures were acquired using a ChemiDoc Imager (BioRad). Antibodies used are listed in Supplemental Table 3 and 4.

### Primary breast organoid Reconstitution Assay

K14-GFP primary tumors were isolated down to the single cell level using sequential collagenase followed by trypsin digestion. Single cells were strained (70 µm) to remove any debris or remaining cell clusters, and stained for live/dead (FxCycle Violet), Cdh3-PE (R&D systems), PDGFR-PE-Cy7 (ThermoFisher), and Thy-1.1-PE-Cy7 (ThermoFisher). FACS sorting (BD Aria II) was performed in a sequential order: first, cells were gated for live cells, second, cells were gated to remove CAFs (PDGFR and Thy-1.1), and finally, cells were gated to sort out K14+/Cdh3+ cells. K14+/Cdh3+ cells were mixed with normal mammary gland (MG) organoids and cultured in low-adhesion well plates (Corning) overnight (37°C, 20% O_2_) to form “MG+K14^+^/Cdh3^+^” organoids. Normal mammary gland organoids were isolated from female wild-type FVB/n age-matched mice and digested to the organoid level using collagenase 1 solution(18, 22).

### Chromatin immunoprecipitation (ChIP) Assay

The Chromatin immunoprecipitation Assay was performed with commercial reagents (Millipore, Cat #17-295). In brief, confluent (cell-cell contacted) WT 4T1 cells were fixed with 1% formaldehyde at 37 °C for 10 min. and stopped with 100 mM glycine for 5min. After washing with PBS, cells were collected and lysed in SDS lysis buffer (1% SDS, 10 mM EDTA, 50 mM Tris, pH 8.1, supplemented with protease inhibitor), then sonicated into 200 to 1,000 bp fragments. Sonicated chromatin was diluted 10-fold with ChIP dilution buffer (0.01% SDS, 1.1% Triton X100, 1.2 mM EDTA, 16.7 mM Tris-HCl, pH 8.1, 167 mM NaCl), then incubated overnight 4°C with the following antibodies: β-catenin (mouse monoclonal, Cat# 7963, Santa Cruz), TCF-4 (mouse monoclonal, Cat# CS204338, Millipore). The normal mouse IgG (Cat# 12-371B, Millipore) was used as a negative control. The next day, 60 µl protein A agarose/Salmon Sperm DNA was added to each sample and incubated at 4°C for 1 hr, followed by four washes with wash buffer. The protein-DNA complexes were eluted with freshly prepared elution buffer (1%SDS, 0.1M NaHCO_3_) and crosslink reversed (5M NaCl, 65°C for 4 hrs) with subsequent addition of proteinase K (45°C for 1 hr). The DNA fragments were purified by phenol/chloroform extraction, followed by ethanol precipitation. Quantitative analysis of the DNA fragments was performed by SYBR Green qPCR. The qPCR data were analyzed by the fold enrichment method using 2^−ΔΔCt^ formula. Primers used are provided in Supplemental Table 6. The SP5 promoter was used as a positive control for β-catenin and TCF4 ChIP.

### Statistical Analysis

For all organoid experiments we analyzed a minimum of 12 organoids from at least 3 different mice in the microfluidic devices. Assuming a 50% change between experimental and control groups, a 0.05 significance level and 0.80 power we estimated that we need at least 10 organoids per genotype per experimental condition. Although most alleles we study are on a pure FVB/n background not all are, therefore, the use of 3 or more different mice controlled for potential phenotype-strain variability. All data was analyzed by t-test and ANOVA with Tukey’s Post-hoc analysis using GraphPad Prism 8.

## Acknowledgments

We thank the following people: Ashley C King, Caleb McCurdy and Jessanne Y Lichtenberg for assistance synthesizing microfluidic devices, Audrey Brenot for feedback on the work, Takako Sasaki for the Laminin α3 antibody (1110+), Prabhakar Andhey and Maxim Artyomov (Washington University) for bioinformatic assistance, Cara Gottardi (Northwestern University) for assistance in design of β-catenin ChIP experiments, the Washington University Center for Cellular Imaging (WUCCI) and VCU Nanomaterials Core Characterization Facility for imaging assistance, the Flow Cytometry and Fluorescence Activated Cell Sorting Core at Washington University for assistance sorting cells, and the McDonnell Genome Institute at Washington University for assistance with single cell sequencing experiments.

## Author contributions

Conceptualization: PYH, AP, GDL

Methodology: PYH, DC, YC, MC, JA

Investigation: PYH, DC, YC, MC, JA

Computational Modeling: JM, AP

Supervision: PYH, AP, GDL

Writing-original draft: PYH, AP, GDL

## Data and materials availability

All data, code, and materials used in the analyses must be available in some form to any researcher for purposes of reproducing or extending the analyses. Include a note explaining any restrictions on materials, such as materials transfer agreements (MTAs). Include accession numbers to any data relevant to the paper and deposited in a public database; include a brief description of the dataset or model with the number. If all data are in the paper and supplementary materials, include the sentence, “All data are available in the main text or the supplementary materials.”

## References

1. Shellard A, Mayor R. Rules of collective migration: from the wildebeest to the neural crest. Philosophical Transactions of the Royal Society B: Biological Sciences. 2020;375(1807):20190387-.

2. Capuana L, Boström A, Etienne-Manneville S. Multicellular scale front-to-rear polarity in collective migration. Current Opinion in Cell Biology. 2020;62:114–22.

3. Shellard A, Mayor R. All Roads Lead to Directional Cell Migration. Trends in Cell Biology. 2020.

4. Aceto N, Toner M, Maheswaran S, Haber DA. En Route to Metastasis: Circulating Tumor Cell Clusters and Epithelial-to-Mesenchymal Transition. Trends in Cancer. 2015;1(1):44–52.

5. Friedl P, Gilmour D. Collective cell migration in morphogenesis, regeneration and cancer. Nature Reviews Molecular Cell Biology. 2009;10(7):445–57.

6. Etienne-Manneville S. Neighborly relations during collective migration. Current Opinion in Cell Biology. 2014;30:51–9.

7. van Helvert S, Storm C, Friedl P. Mechanoreciprocity in cell migration. Nature cell biology. 2018;20(1):8–20.

8. Ewald AJ, Brenot A, Duong M, Chan BS, Werb Z. Collective epithelial migration and cell rearrangements drive mammary branching morphogenesis. Dev Cell. 2008;14(4):570–81.

9. Khalil AA, Friedl P. Determinants of leader cells in collective cell migration. Integr Biol (Camb). 2010;2(11-12):568–74.

10. Mayor R, Etienne-Manneville S. The front and rear of collective cell migration. Nat Rev Mol Cell Biol. 2016;17(2):97–109.

11. Chen BJ, Tang YJ, Tang YL, Liang XH. What makes cells move: Requirements and obstacles for leader cells in collective invasion. Exp Cell Res. 2019;382(2):111481-.

12. Vishwakarma M, Di Russo J, Probst D, Schwarz US, Das T, Spatz JP. Mechanical interactions among followers determine the emergence of leaders in migrating epithelial cell collectives. Nature Communications. 2018;9(1):3469-.

13. Aceto N, Bardia A, Miyamoto DT, Donaldson MC, Wittner BS, Spencer JA, et al. Circulating tumor cell clusters are oligoclonal precursors of breast cancer metastasis. Cell. 2014;158(5):1110–22.

14. Bulfoni M, Turetta M, Del Ben F, Di Loreto C, Beltrami AP, Cesselli D. Dissecting the Heterogeneity of Circulating Tumor Cells in Metastatic Breast Cancer: Going Far Beyond the Needle in the Haystack. International journal of molecular sciences. 2016;17(10):1775-.

15. Au SH, Storey BD, Moore JC, Tang Q, Chen YL, Javaid S, et al. Clusters of circulating tumor cells traverse capillary-sized vessels. Proc Natl Acad Sci U S A. 2016;113(18):4947–52.

16. Allen TA, Asad D, Amu E, Hensley MT, Cores J, Vandergriff A, et al. Circulating tumor cells exit circulation while maintaining multicellularity, augmenting metastatic potential. Journal of Cell Science. 2019;132(17):jcs231563–jcs.

17. Nguyen-Ngoc K-V, Cheung KJ, Brenot A, Shamir ER, Gray RS, Hines WC, et al. ECM microenvironment regulates collective migration and local dissemination in normal and malignant mammary epithelium. Proceedings of the National Academy of Sciences. 2012;109(39):E2595–E604.

18. Cheung Kevin J, Gabrielson E, Werb Z, Ewald Andrew J. Collective Invasion in Breast Cancer Requires a Conserved Basal Epithelial Program. Cell.155(7):1639–51.

19. Cheung KJ, Padmanaban V, Silvestri V, Schipper K, Cohen JD, Fairchild AN, et al. Polyclonal breast cancer metastases arise from collective dissemination of keratin 14-expressing tumor cell clusters. Proc Natl Acad Sci U S A. 2016;113(7):E854–63.

20. Cheung KJ, Ewald AJ. A collective route to metastasis: Seeding by tumor cell clusters. Science. 2016;352(6282):167–9.

21. Theveneau E, Linker C. Leaders in collective migration: are front cells really endowed with a particular set of skills? F1000Res. 2017;6:1899.

22. Hwang PY, Brenot A, King AC, Longmore GD, George SC. Randomly distributed K14<sup>þ</sup> breast tumor cells polarize to the leading edge and guide collective migration in response to chemical and mechanical environmental cues. Cancer Research. 2019;79(8).

23. Sachs N, Clevers H. Organoid cultures for the analysis of cancer phenotypes. Current opinion in genetics & development. 2014;24:68–73.

24. Drost J, Clevers H. Organoids in cancer research. Nature reviews Cancer. 2018;18(7):407–18.

25. Lawson DA, Kessenbrock K, Davis RT, Pervolarakis N, Werb Z. Tumour heterogeneity and metastasis at single-cell resolution. Nature cell biology. 2018;20(12):1349–60.

26. Turashvili G, Brogi E. Tumor Heterogeneity in Breast Cancer. Frontiers in Medicine. 2017;4(227).

27. Brouwer A, De Laere B, Peeters D, Peeters M, Salgado R, Dirix L, et al. Evaluation and consequences of heterogeneity in the circulating tumor cell compartment. Oncotarget. 2016;7(30):48625–43.

28. Miyamoto DT, Ting DT, Toner M, Maheswaran S, Haber DA. Single-Cell Analysis of Circulating Tumor Cells as a Window into Tumor Heterogeneity. Cold Spring Harbor symposia on quantitative biology. 2016;81:269–74.

29. Benyahia Z, Dussault N, Cayol M, Sigaud R, Berenguer-Daizé C, Delfino C, et al. Stromal fibroblasts present in breast carcinomas promote tumor growth and angiogenesis through adrenomedullin secretion. Oncotarget. 2017;8(9).

30. Toullec A, Gerald D, Despouy G, Bourachot B, Cardon M, Lefort S, et al. Oxidative stress promotes myofibroblast differentiation and tumour spreading. EMBO Molecular Medicine. 2010;2(6):211–30.

31. Katt ME, Placone AL, Wong AD, Xu ZS, Searson PC. In Vitro Tumor Models: Advantages, Disadvantages, Variables, and Selecting the Right Platform. Frontiers in Bioengineering and Biotechnology. 2016;4.

32. Weber CE, Kuo PC. The tumor microenvironment. Surgical Oncology. 2012;21(3):172–7.

33. Clark AG, Vignjevic DM. Modes of cancer cell invasion and the role of the microenvironment. Current Opinion in Cell Biology. 2015;36:13–22.

34. Emon B, Bauer J, Jain Y, Jung B, Saif T. Biophysics of Tumor Microenvironment and Cancer Metastasis -A Mini Review. Computational and Structural Biotechnology Journal. 2018;16:279–87.

35. Bartoschek M, Oskolkov N, Bocci M, Lovrot J, Larsson C, Sommarin M, et al. Spatially and functionally distinct subclasses of breast cancer-associated fibroblasts revealed by single cell RNA sequencing. Nat Commun. 2018;9(1):5150.

36. Corsa CA, Brenot A, Grither WR, Van Hove S, Loza AJ, Zhang K, et al. The Action of Discoidin Domain Receptor 2 in Basal Tumor Cells and Stromal Cancer-Associated Fibroblasts Is Critical for Breast Cancer Metastasis. Cell Rep. 2016;15(11):2510–23.

37. Sridhar S, Rajesh C, Jishnu PV, Jayaram P, Kabekkodu SP. Increased expression of P-cadherin is an indicator of poor prognosis in breast cancer: a systematic review and meta-analysis. Breast Cancer Res Treat. 2020;179(2):301–13.

38. Vieira AF, Dionísio MR, Gomes M, Cameselle-Teijeiro JF, Lacerda M, Amendoeira I, et al. P-cadherin: a useful biomarker for axillary-based breast cancer decisions in the clinical practice. Mod Pathol. 2017;30(5):698–709.

39. Plutoni C, Bazellieres E, Le Borgne-Rochet M, Comunale F, Brugues A, Seveno M, et al. P-cadherin promotes collective cell migration via a Cdc42-mediated increase in mechanical forces. J Cell Biol. 2016;212(2):199–217.

40. Franzke CW, Tasanen K, Schäcke H, Zhou Z, Tryggvason K, Mauch C, et al. Transmembrane collagen XVII, an epithelial adhesion protein, is shed from the cell surface by ADAMs. Embo j. 2002;21(19):5026–35.

41. Bayer SV, Grither WR, Brenot A, Hwang PY, Barcus CE, Ernst M, et al. DDR2 controls breast tumor stiffness and metastasis by regulating integrin mediated mechanotransduction in CAFs. Elife. 2019;8.

42. Qian M, Wang DC, Chen H, Cheng Y. Detection of single cell heterogeneity in cancer. Semin Cell Dev Biol. 2017;64:143–9.

43. Shen Y, Schmidt BUS, Kubitschke H, Morawetz EW, Wolf B, Käs JA, et al. Detecting heterogeneity in and between breast cancer cell lines. Cancer Converg. 2020;4(1):1.

44. Pearson A, Smyth E, Babina IS, Herrera-Abreu MT, Tarazona N, Peckitt C, et al. High-Level Clonal FGFR Amplification and Response to FGFR Inhibition in a Translational Clinical Trial. Cancer Discov. 2016;6(8):838–51.

45. Sethi N, Dai X, Winter CG, Kang Y. Tumor-derived JAGGED1 promotes osteolytic bone metastasis of breast cancer by engaging notch signaling in bone cells. Cancer Cell. 2011;19(2):192–205.

46. Bednarz-Knoll N, Efstathiou A, Gotzhein F, Wikman H, Mueller V, Kang Y, et al. Potential Involvement of Jagged1 in Metastatic Progression of Human Breast Carcinomas. Clin Chem. 2016;62(2):378–86.

47. Xing F, Okuda H, Watabe M, Kobayashi A, Pai SK, Liu W, et al. Hypoxia-induced Jagged2 promotes breast cancer metastasis and self-renewal of cancer stem-like cells. Oncogene. 2011;30(39):4075–86.

48. Idoux-Gillet Y, Nassour M, Lakis E, Bonini F, Theillet C, Du Manoir S, et al. Slug/Pcad pathway controls epithelial cell dynamics in mammary gland and breast carcinoma. Oncogene. 2018;37(5):578–88.

49. Khalil AA, de Rooij J. Cadherin mechanotransduction in leader-follower cell specification during collective migration. Exp Cell Res. 2019;376(1):86–91.

50. Labernadie A, Kato T, Brugues A, Serra-Picamal X, Derzsi S, Arwert E, et al. A mechanically active heterotypic E-cadherin/N-cadherin adhesion enables fibroblasts to drive cancer cell invasion. Nature cell biology. 2017;19(3):224–37.

51. Christgen M, Bartels S, van Luttikhuizen JL, Bublitz J, Rieger LU, Christgen H, et al. E-cadherin to P-cadherin switching in lobular breast cancer with tubular elements. Modern Pathology. 2020.

52. Albergaria A, Ribeiro A-S, Vieira A-F, Sousa B, Nobre A-R, Seruca R, et al. P-cadherin role in normal breast development and cancer. The International Journal of Developmental Biology. 2011;55(7-8-9):811–22.

53. Baohua Li Fenfen Wang XWDG. P-cadherin promotes cervical cancer growth and invasion through affecting the expression of E-cadherin and p120 catenin. European Journal of Gynaecological Oncology.40(2):224–31.

54. Imai S, Kobayashi M, Takasaki C, Ishibashi H, Okubo K. High expression of P-cadherin is significantly associated with poor prognosis in patients with non-small-cell lung cancer. Lung Cancer. 2018;118:13–9.

55. Jasaitis A, Estevez M, Heysch J, Ladoux B, Dufour S. E-cadherin-dependent stimulation of traction force at focal adhesions via the Src and PI3K signaling pathways. Biophys J. 2012;103(2):175–84.

56. Mertz AF, Che Y, Banerjee S, Goldstein JM, Rosowski KA, Revilla SF, et al. Cadherin-based intercellular adhesions organize epithelial cell-matrix traction forces. Proc Natl Acad Sci U S A. 2013;110(3):842–7.

57. Le Borgne-Rochet M, Angevin L, Bazellières E, Ordas L, Comunale F, Denisov EV, et al. P-cadherin-induced decorin secretion is required for collagen fiber alignment and directional collective cell migration. J Cell Sci. 2019;132(21).

58. Vieira AF, Ribeiro AS, Dionisio MR, Sousa B, Nobre AR, Albergaria A, et al. P-cadherin signals through the laminin receptor alpha6beta4 integrin to induce stem cell and invasive properties in basal-like breast cancer cells. Oncotarget. 2014;5(3):679–92.

59. Walko G, Castanon MJ, Wiche G. Molecular architecture and function of the hemidesmosome. Cell Tissue Res. 2015;360(3):529–44.

60. Sanghvi-Shah R, Weber GF. Intermediate Filaments at the Junction of Mechanotransduction, Migration, and Development. Front Cell Dev Biol. 2017;5:81.

61. Wang W, Zuidema A, Te Molder L, Nahidiazar L, Hoekman L, Schmidt T, et al. Hemidesmosomes modulate force generation via focal adhesions. J Cell Biol. 2020;219(2).

62. Bonakdar N, Schilling A, Sporrer M, Lennert P, Mainka A, Winter L, et al. Determining the mechanical properties of plectin in mouse myoblasts and keratinocytes. Exp Cell Res. 2015;331(2):331–7.

63. Na S, Chowdhury F, Tay B, Ouyang M, Gregor M, Wang Y, et al. Plectin contributes to mechanical properties of living cells. Am J Physiol Cell Physiol. 2009;296(4):C868–77.

64. Swat MH, Thomas GL, Belmonte JM, Shirinifard A, Hmeljak D, Glazier JA. Multi-scale modeling of tissues using CompuCell3D. Methods Cell Biol. 2012;110:325–66.

65. Vaezi A, Bauer C, Vasioukhin V, Fuchs E. Actin cable dynamics and Rho/Rock orchestrate a polarized cytoskeletal architecture in the early steps of assembling a stratified epithelium. Dev Cell. 2002;3(3):367–81.

66. Costa A, Kieffer Y, Scholer-Dahirel A, Pelon F, Bourachot B, Cardon M, et al. Fibroblast Heterogeneity and Immunosuppressive Environment in Human Breast Cancer. Cancer Cell. 2018;33(3):463–79.e10.

67. Wagner J, Rapsomaniki MA, Chevrier S, Anzeneder T, Langwieder C, Dykgers A, et al. A Single-Cell Atlas of the Tumor and Immune Ecosystem of Human Breast Cancer. Cell. 2019;177(5):1330–45.e18.

