## Supplementary Materials for "A Cdh3-Lam332 signaling axis in a leader cell subpopulation controls protrusion dynamics and tumor organoid collective migration"

**This PDF file includes:**

Figs. S1 to S7
Tables S1 to S6

**Fig. S1.**

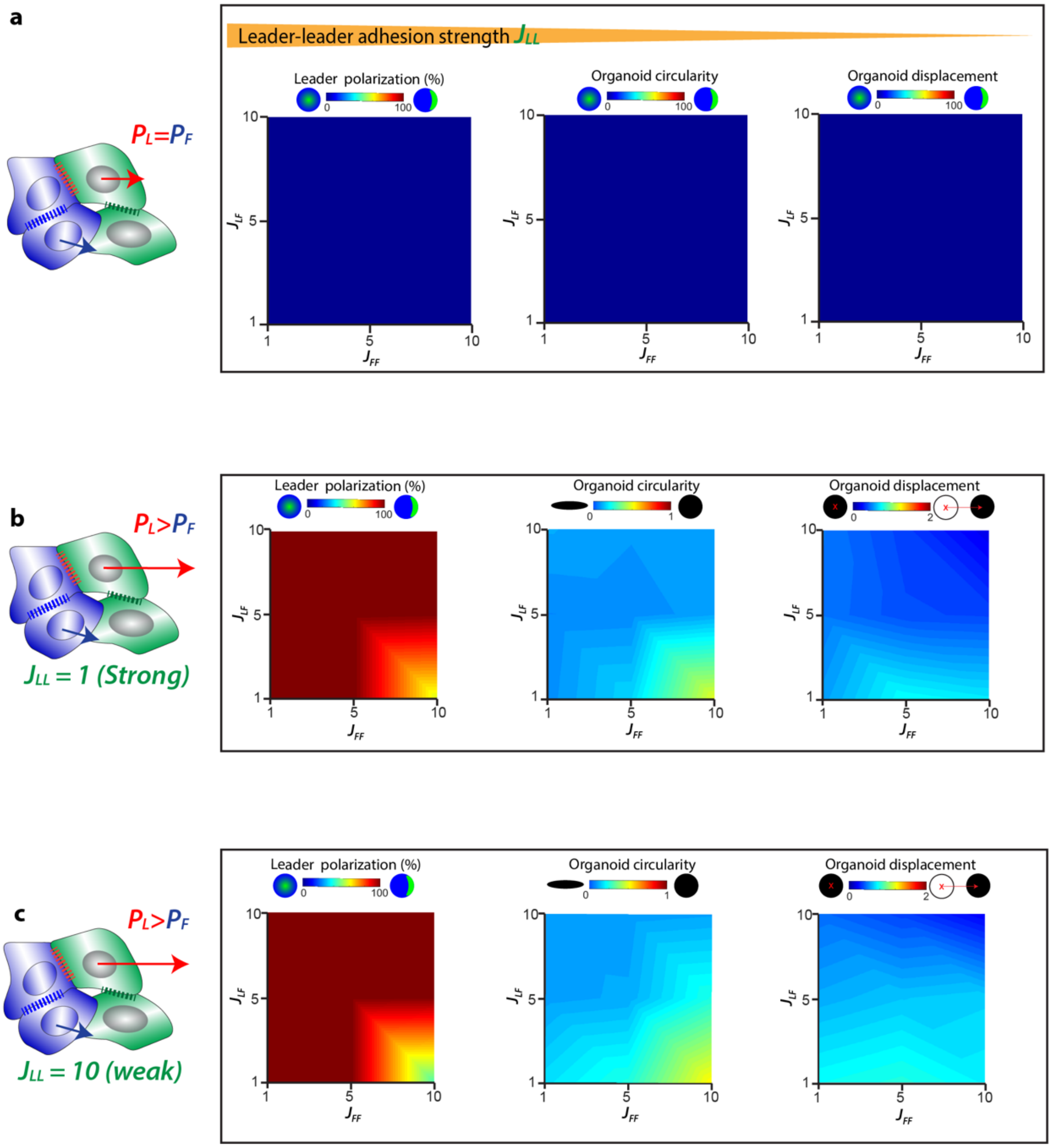


**Fig. S1: Simulation results showing the effect of differential adhesions and protrusions on organoid migration.** After simulations are performed in a parametric space varying leader-follower adhesion ($J_{LF}$) and follower-follower adhesion ($J_{FF}$), leader polarization, organoid circularity, and organoid displacement are plotted for these cases: **(a)** when leader and follower protrusions are the same, leaders do not polarize and organoid does not move regardless of any possible combination of leader/follower adhesions. **(b, c)** When leader protrusions are higher than followers (which are all other cells in the organoids), leader cells polarization increases but net organoid displacement remains low regardless of any tested leader/follower adhesion combinations.

**Fig. S2.**

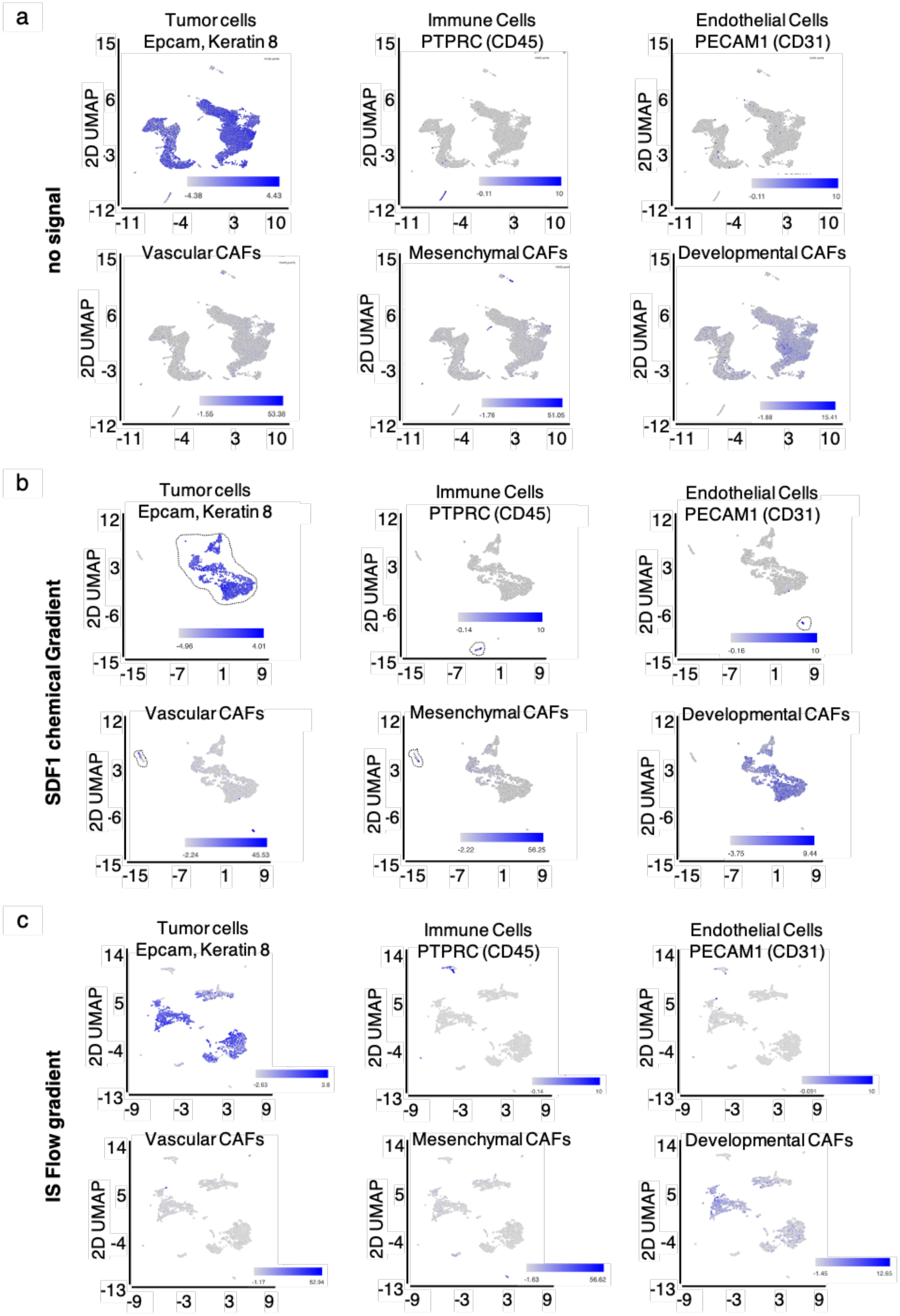


**Fig. S2: scRNA-seq distinguishes major cell types within primary tumor organoids.** Markers used to identify unique cell types that identify clusters for (a) no signal, (b) SDF1 chemical gradient, and (c) IS Flow gradient.

**Fig. S3.**

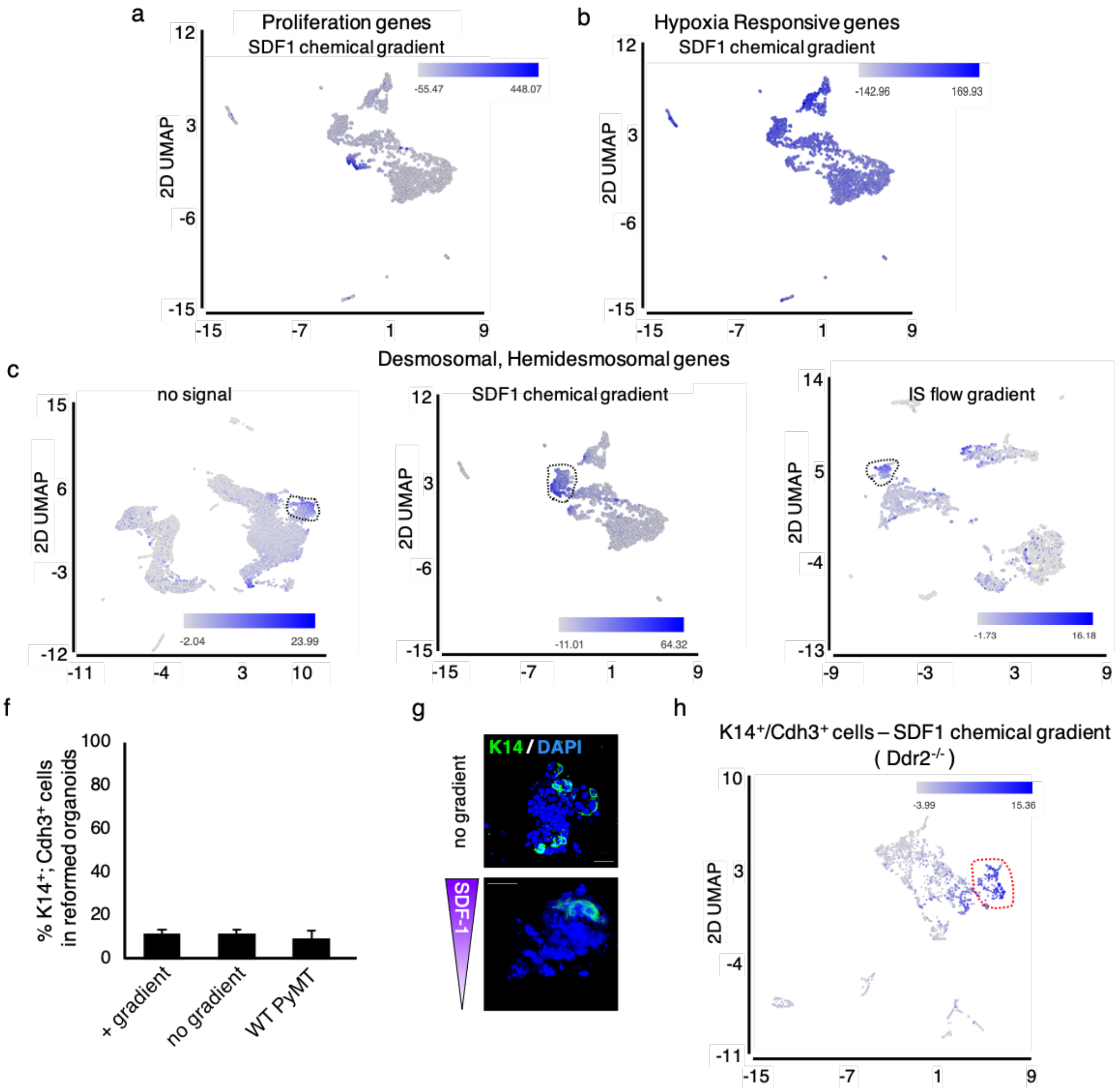


**Fig. S3: Characterization of K14+ leader cell subsets and organoid reconstitution assay. (a and b)** UMAP plots of organoids exposed to a SDF1 chemical gradient showing cluster 6 expresses cell proliferation genes (a) and clusters with enriched hypoxia responsive genes (b). **(c-e)** UMAP plots showing clusters enriched for desmosomal, hemidesmosomal genes for no signal, SDF1 chemical gradient, and IS Flow gradient. **(f)** Quantification of percent of K14+;Cdh3+ leader cells in reformed organoids before and after exposure to SDF1 gradient, compared to WT PyMT organoids. **(g)** Representative IF images of K14 expression in reconstituted organoids before and after SDF1 chemical gradient. **(h)** UMAP plot showing K14+;Cdh3+ cluster in MMTV-PyMT DDR2-/- organoids after exposure to a SDF1 chemical gradient.

**Fig. S4.**

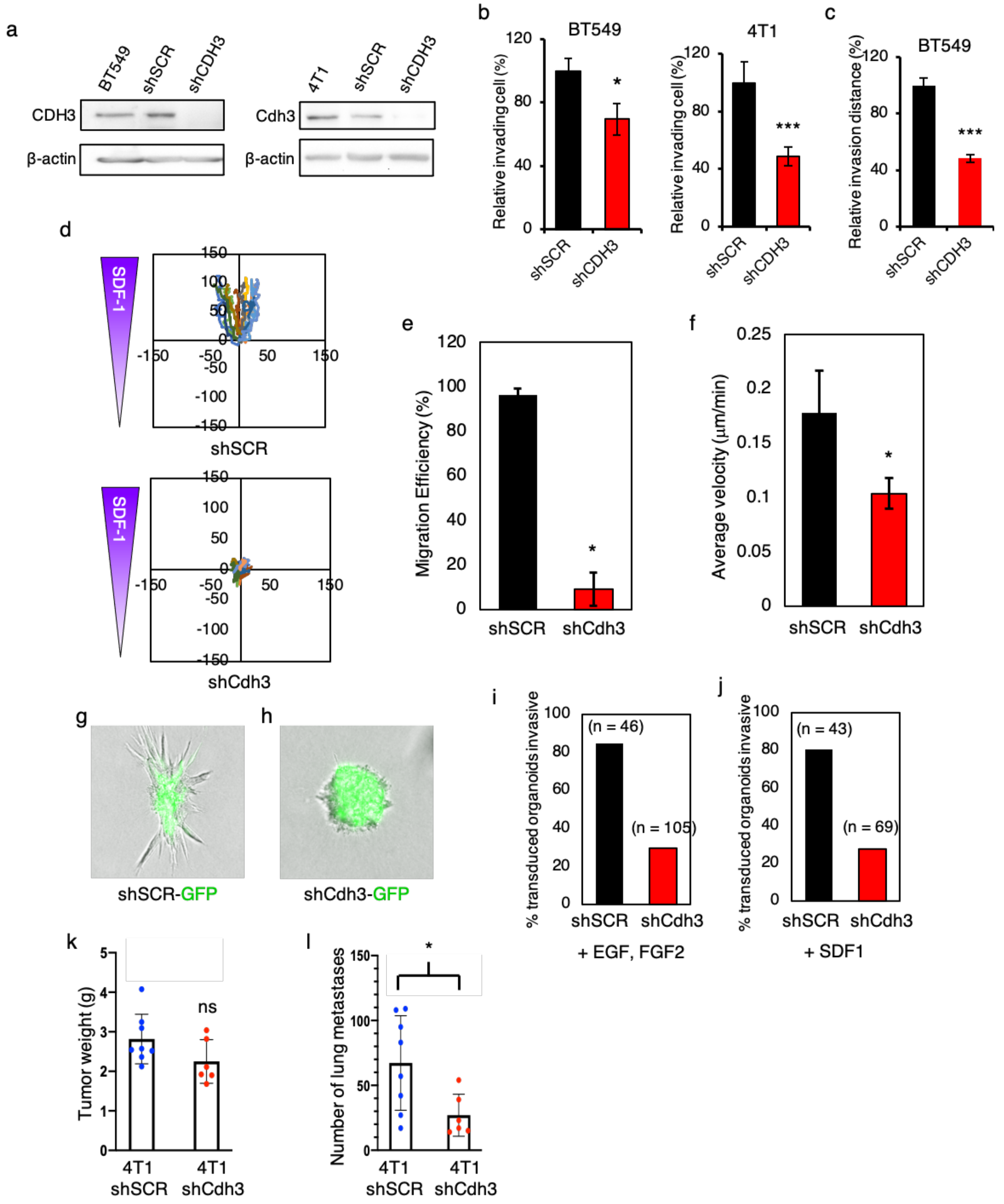


**Fig. S4: Cadherin 3 expression is required for leader cell function. (a)** Western blot verification shCdh3 lentiviral depletion in human BT549 and mouse 4T1 breast tumor cells. **(b)** Percent of BT549 and 4T1cells (+/- Cdh3) invading through matrigel-coated transwells. **(c)** Relative invasion of BT549 cells (+/- Cdh3) through collagen I coated transwells. **(d)** Collective migration tracking maps for shCdh3 and control shSCR transduced human PDX breast tumor organoids after exposure to SDF1 gradients. (**e and f**) Quantification of human breast tumor PDX organoid migration efficiency and average velocity in the direction of the SDF1 gradient. **(g and h)** Representative IF images of invasive MMTV-PyMT primary organoids transduced with shCdh3-GFP or control shSCR-GFP lentiviruses cultured in standard 3D collagen I gels, normoxia, and uniform concentration of added cytokines. **(i and j)** Quantification of invasive organoids following transduction with shCDH3-GFP or control shSCR-GFP lentiviruses and cultured in standard 3D collagen I, normoxia and uniform cytokine concentration: EGF + FGF2 **(i)** or SDF1 **(j).** **(k and l)** Cdh3 depletion results in decreased lung metastasis in 4T1 orthotopic breast cancer mouse model. **(k)** Tumor weights. **(l)** Number of total lung metastases (by histologic counting) after primary tumors (4T1 shSCR and 4T1 shCdh3) reached endpoint (@2 cm). For all experiments, $^*p<0.05$, $^{***}p<0.001$ ANOVA with Tukey's post hoc analysis.

**Fig. S5.**

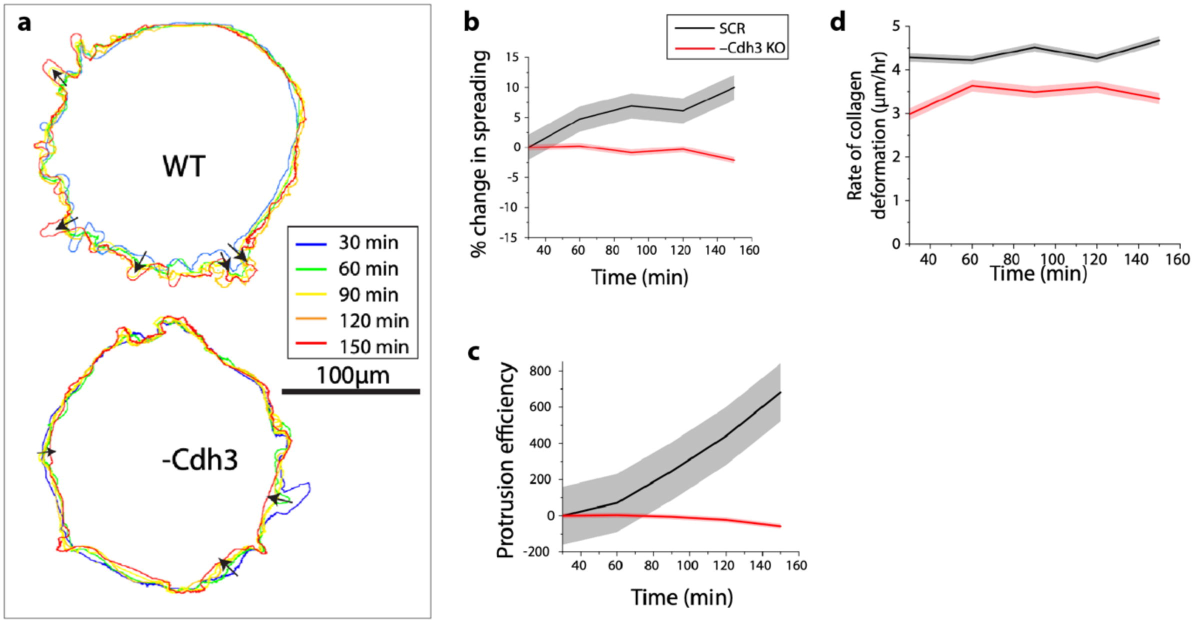


**Fig. S5: Cdh3 depletion in BT549 cells reduces protrusion efficiency and collagen deformation. (a)** Human BT-549 cells control (shSCR) and Cdh3-knockdown (shCdh3) are constituted as spheroids from approximately 5000-8000 cells, encased in 2.3 mg/ml collagen for 18h, and live imaging performed. Protrusion outlines for representative BT549 spheroids are plotted to demonstrate protrusion dynamics over time, visualize by the color-coded timestamps. Scale bar = 100 µm. Arrows denote unidirectional (single arrowheads) and regressive (double arrowheads) protrusions in shSCR and shCdh3 cases, respectively. **(b)** Percentage change in organoid area, relative to the reference timepoint, over time calculated from protrusion outlines. **(c)** Protrusion efficiency (%area-min) calculated in terms of area under the curve of percent spreading over time using Y=0 as baseline; here, regressive protrusions reduce protrusion efficiency. **(d)** Rate of collagen deformation of live spheroids (before trypsinization) over time. N=3 for shSCR and N=3 for shCdh3. In plots, solid lines represent mean and shaded background represents standard error (SE).

**Fig. S6.**

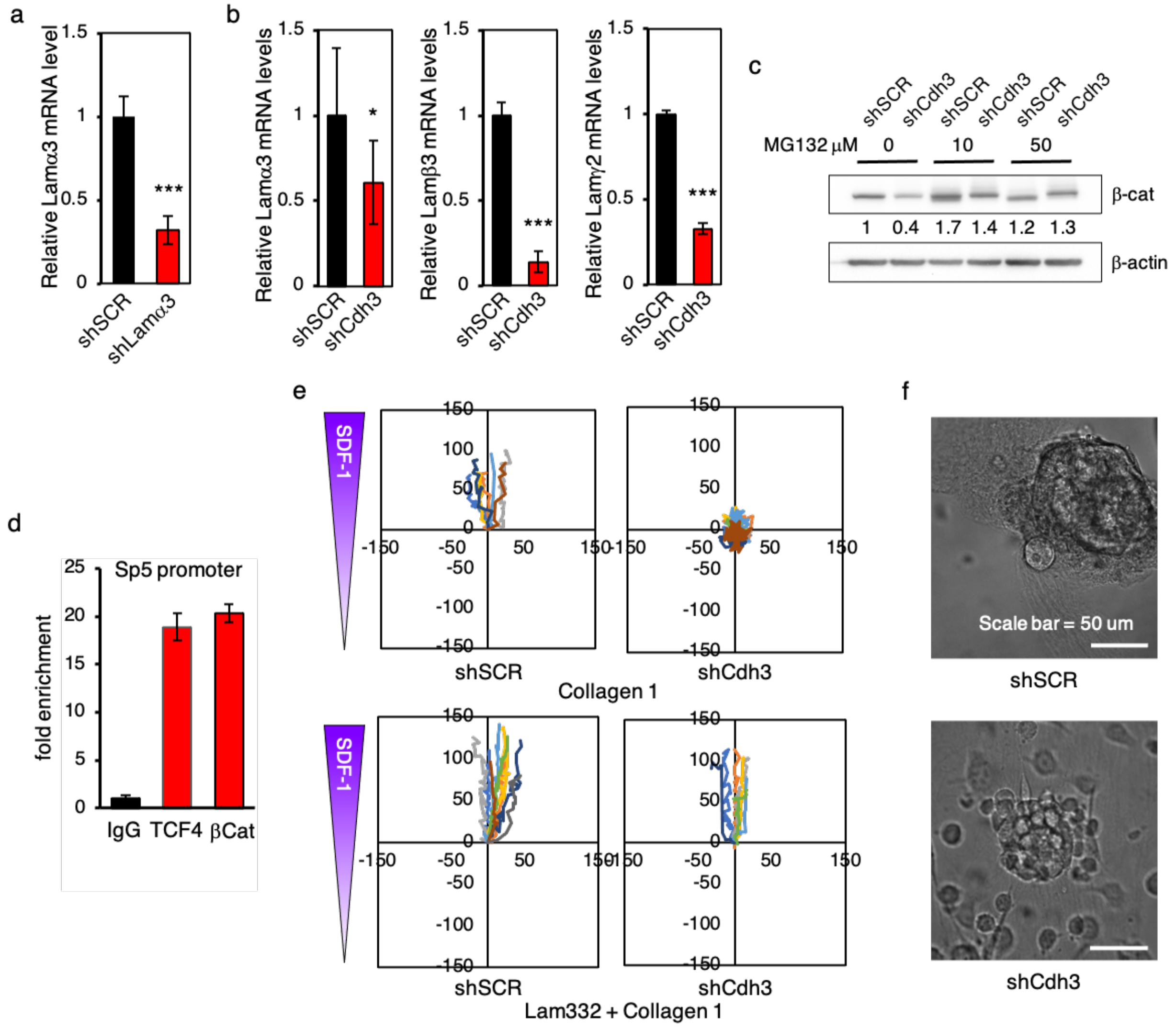


**Fig. S6: Cadherin 3 regulates ECM feedback through control of Laminin 332 expression by leader cells. (a)** Efficacy of Laminin a3 shRNA lentiviruses depletion of Laminin a3 in mouse 4T1 cells: Q-PCR. **(b)** Laminin a3, b3, and g2 mRNA expression in aggregated 4T1 cells depleted of Cadherin 3 (shCdh3). **(c)** Proteosome inhibition stabilizes b-catenin in aggregated 4T1 cells lacking Cdh3 expression. Western blot with the indicated antibodies of cell lysates from aggregated control (shSCR) or Cadherin 3 depleted (shCdh3) 4T1 cells treated, or not, with the indicated doses of MG132 for 5 h. **(d)** Positive control chromatin IP (IgG, TCF4, b-Catenin antibodies) using PCR primers for the Sp5 promoter. **(e)** Cluster motility tracking of control (shSCR) or Cdh3-depleted (shCdh3) aggregated 4T1 cell clusters placed in microfluidic devices in either 3D Collagen I (upper) or 3D Collagen I:Laminin 332 (1:1) (lower) and hypoxia and an SDF1 gradient. **(f)** Images of control (shSCR) and Cdh3-depleted (shCdh3) 4T1 spheroids in a 3D collagen I/laminin 332 matrix within microfluidic devices.

**Fig. S7.**

putative Laminin α3 promoter

```
-1505  AGCCCTGGCTGTCCTGAAACTCGCTCCGTAAACCAGGCTGGCCTCAAACTCAAGAGATCCTCCTGCCTCT
-1435  GCCTTTGGAGTGCTGGGATTAAAGGTGTGTGGCACTGTTGCTGAAGGGCTCATTTGAGTATTTACAGCAG
-1365  AGTAACACAAAGTAACACAAAGCCTGCAAACACAAAATTGTGTGAAATCAGAACAAGGTGTCAGCAGATA
-1295  TCGTTTGTGGTGGGGGCTGCTTCCTGGTTTAGACAGTCTTCACTGTGCCCTTATAGTAGCGGGAGCCAGC
-1225  CAGCCAGCACACTTTCTCTCTCTCTCTCTCTCTCTCTCTCTCTCTCTCTCTCTCTCTCTCTCTCTCTCTC
-1155  TCTCCATTCATTTATTCATTTTGGTTTGAGATAGGGTTTTGTCAGCCCAGGCTGATCTTCGACACAGGAC
-1085  GCACCTGCCTCAGCCTTCCATAGTGCTGGGTTGCAGGTGTGTGCCGCCATGCCCTTGGGGCTCGATTATA
-1015  AAGCCACTGATCACGTTCACCGGTACCACCCCCTCGCTGTATTTATTGATTGCTCCCTGGATATAAATCT
 -945  GTACAAGTGGGATGACTGAAGCAAATGCATACTAGAGTTATTTACCAAGAAACGAAAAAAAAAATCCTTT
 -875  AACTGAAACCTTCCTGGTTGATCATAGGGTTTCAGCCTATGAATTTGTTGGGGGGAGGTTCAGGGATATT
 -805  CACACCACAGCACCTCTCAGGTGTACCAAGTGTTCTCATTTCTGTATCCAGGCCGAGCAGTGTTCTCACG
 -735  TCTTGCCTCTTTCCCCTCCTCTAGCCTTCAAGGCTTCAGTTTCATCTGGGATACTTGGAGCCCTTGAGCA
 -665  TGCTGGCTTCCCCATCTGTGGCACCTCCTCTAATAGCTTAGAGGGCCTACTGTAGACTCAATGTTGGAGG
 -595  ATGTCCAAGCCTCAGCCTTAAAGGAGTAATACCGGGACATCTAAAGGATGTATTTCCCCAAGGTACCTTC
 -525  TCTGCTAACCACGGAGTGCACCATGGGTTGTGGTTTGAATGTATACTTTTCTGCCACTGCTAATTCAGCA
 -455  AGATCCAGACTCTGTACGTAGCTGTAGCGGCAAGGTAGAAGAATGGGCCAACAGATCAACGCTCGCCTGG
 -385  GAACTGGAAACGCCTTTGAGGAAAGTTCTACAGGCAGGGGCAGTTTCCTTCAAGCTTCTCAGGAGCTGGA
 -315  GTATTTGGAGAGCTTGATTTCTGACCTGGCTCTTAGAACAGTCAGAGGAACTCCCCTGGTCTTCTGGATT
 -245  TGGGGAGCCCGGGAAAGTTTGCAGCCAGCAATGTCCACGCCTGACATTTCCTGATCCCGAGGGTGCCAGC
 -175  AACAGCGGCAGGTTCTCAACCCGGGGCCACGCCCACGCGGCGGGCTCATCGCGTGGACAGAGGGGAGCGG
 -105  CCCAGGGCGCCGCCCACTCCCGGGGCTGTCCAGGCGTGCTGAGAGTAGCAGCCTCCTGCTCCGCGGTGCT
  -35  ACCCTGGGCACAGGCAGGAGACCCAGCGCCGCGGG@ATGGCGGTGGCTCTGGGCAGGGCACCGCGATCCT
```

putative Laminin β3 promoter

```
-1198  GCTAAAATCTGGGCTTAGTACGAAGGCAAGGTGGGAGGAATGGTCATGGCCGGGCTCTGCTCAGCCTCAT
-1128  TCTTGGTGGGTTTTCAGCTGGGGCATGAGTGCAGAGAGTACTGCGAAATGTCAGGGATGCTTTCTGTCTT
-1058  TGACCCTCAGTGGCCCTGAGACCTCAGGCCACAGTCCCAGCTTGGGTTCCCGACTGCTCTGTAGGAAGGA
 -988  CATCCATCGAAGCATTGGGATGTAATGGCAGAAAGGCTGCTGGTTCCCCTCTGCCTGACCACCATCCATC
 -918  TTTCTTCCCCAGCTGCTTGGATGTCATTAGGCTGACCAGTGGAGGAAGGGTGGTGCATGCCTCTGAAGTG
 -848  GGCCTGCCTCCTGCACTGTTTCCAGCTGCCTTCTGTGCTTTTGTTCAGAACGGGAGATGCTTTGAGATGC
 -778  TCTTACTTTTCTTAAAACTGTCGGGTCCACCTCTCCCCACAGCACCCCACCTCTCCCCACAGCACCCCAC
 -708  CTCTCCCCACGGCACCCCACCTTTCCCCACAGCACCCCACCTCTCCCCACGGCACCCCACCTTTCCCCAC
 -638  AGCACCCTACCTCTCCCCACAGCACCTCCCCCCTCCCCACAGCACCCATGTTTCCCACAGTGCCACCTC
 -568  ACCTCTCCCCAAGGCACCCCAGGTGGCTTTTAAAGATCTCTCTTCTTCCCATAGCATTCCTGCAGCCTGG
 -498  TTTTCCTAGAATTGTGGCTACCACAGCTTCACCTGCACTGTAGCCTTGTAATCCTGGAGACCCTGTCAAC
 -428  TCAGGAAATGACCTGCCTCTAGACTTTCAGCACCTTTGGCTTGGGTGCTTTTGGCTTGTACCTAAGCCTG
 -358  GTTTGAAGCCCTGAGAAGCCTGTCCCCCCCCCCCCCCTCTGCAGAAACAAAGCCTTAGACCTCCCTTTAG
 -288  TTATCTGCTGCAATGGGATGTCCTCCCTCCTCCTGCACAGTCCCCATGTTTCCACACAGTTCAGAAGAGG
 -218  AAGCTGATCCTGGGATTTCCCCAGGGCTCCCTCGCAGGTTCTTGCCATTGGCTACAAATGTGCCCACTGC
 -148  TCCAAAGTGTCTTTGTGAAGATCCAGGATGAGGACCTGGTGAGGACAGTGGTCTAGCACAATGAAGAAAG
  -78  GGTGGGGAGGGAGCTTGACCATCCAGGATGCTTTGCTTTGTTCTAGGCGACCCCTCTGAGGACCTGCACT
   -8  GGCTCAGG@ATGACAGCATTTTTCCTCCTGTGGCTTGGTAAGCATCCTTTCCTTCCTCCCATCTCTCTGT
```

putative Laminin γ2 promoter

```
-1478  AGGTTAAGAAAATTAGAATATCCATTCTGTTGCTCATTTTGCGCTAGATTTTAGAGATGTCGGCTTGTCT
-1408  GCAGTACCCTATCTTGCCTCTGTTGCTCTTCAATGCTTTGTCTACTTCCTGGTTAGAACTAGGTATCTAG
-1338  TTCAGACTCCATGTTCTGTACAAAGCCAGCCACATCTCTCCTGGTCAGCCTACACTGGTCACTATCACAC
-1268  TTAGAAATGGGAGTTCTCACACCCTTTGACAATATGTATGGATTACGTTGTTAGTTATCTTAGAGTGTTA
-1198  AGCTACGGAACTATCTATGGTACAGTCATTCATTTAGTCAGTGACTAGGTATTTATTTAGGGCCTCTGCT
-1128  GTCCTTACCATGAGGAAAAGAAGTGGTTATTGGTTGTAATGGATACGTTGTGTCTACATAAATACTAGTA
-1058  AATCCTAAGTCTATAGCAGGATTTAAGAGCCTTCTCTGACAGTCACTTCTGCATTATTCACTGTAGGAGA
 -988  AGTTCCCAGGACAGTGTGGGATAAGCCAAGATTTGCATCCCCTCCCCCACTTGGTGTTACTCACTTCAGC
 -918  TAACACCGGGCTTGGCTGGCACTATCCCTCTTAACGTTATTTTCTAGCGGGGCCAGAGAGAATGTCGTGA
 -848  AAAATATGCTCTGAGATGTTAAAAGACTAAGAAACCTTTTCTCTTATGAGCAGAAGGCTTGTTGAATATC
 -778  AACTGTGAACTAGTTCGTAACAGTTTAATGAGTTTGCTTAATGAGTTTGCTAGTTCAAGGTTAAGGGGGT
 -708  AGATAGACCCAGGAGAGCAACCTTTGGGTCACCCTGGTGTGTGTGTGTGTGTGTGTGTGTGTGTGTGTGT
 -638  GTGTGTGTGTGCTTAGAAGTGAGTAAAATCAGAGGTATGTATACGTGCTGTATCTCAGAATAAAAGTTAC
 -568  CCTGAGAGTGAAGAGGATCGATCCTGTTGGGCCCCTCAGCCTGGTTAATTTGCCTGTGTTTTAGACTAGA
 -498  GTGGAAGGGGCGTCCAACTGTGACTGGAATGGATGTCTGTGCCCTTGAGACTCCCTCGCCCCCTCCGGTT
 -428  TTGAGGAAGGGGTGGCTGATATCTACAACCTGAGTTCTGCCCAGGACCACCCCGGGAGAGAACGGCTGTG
 -358  TGTAGCGGGGTGCGGGTGGAGGGAGGGGATATGGCACATTCCAGAAGGATCTCTGGACCGCAGCCTCAGT
 -288  ACCACGCCCTCGGCGGGGCCAGGTGTGCTCCCCGGTTGCGGAGGGGGGAGCCCGGGGCCGCAGCGAGCGG
 -218  GTTTACTGCTCGGCATGCGCTGAGTGTTTGTCTGCCTCGGAGGGTTGGGTCCTCCTTGTTCACAGGTGAG
 -148  TCACGCCCCGAAACACCGGCTCTCTTCCTGTCAGGACTGAGTCAGGGAGAAGAGTCGATAAAACACCTGA
  -78  TCGAGGGAAAAGGAAGGCACAGCAGAGCGAAGACTCGGCGCCCGGCAGGCACCCGCAGCGGCGGAGCGCG
   -8  GCCCGGCC@ATGCCTGCGCTCTGGCTCAGCTGCTGCCTTGGTGTCGCGCTCCTGCTGCCCGCAGCCCAGG
```

**Fig. S7: Putative promoters of mouse Laminin 332 genes**. Approximately 1500 nucleotides 5' of the ATG start codon for mouse Laminin α3 **(a)**, Laminin β3 (**b)**, and Laminin γ2 **(c)** genes. Highlighted in yellow are potential TCF binding sites within each promoter region. Red arrows identify PCR primers used, for each gene, to amplify β-catenin and TCF4 immunoprecipitated chromatin pieces.

**Table S1.**

Table of all genes identified from scRNA sequencing analysis in K14+/Cdh3+ clusters for no gradient, SDF1, and IS flow is uploaded in the form of "Suppl. Excel" under Auxiliary Supplementary Materials.

**Table S2.**

| shRNA name | Target sequence |
|---|---|
| Laminin $\alpha$3 mouse | 5’ -ATCCAGTGGCTGGCAATATAA- 3’ |
| Cadherin 3 mouse | 5’ -TGAGGACGATGCTGTCAACACTTACAATG- 3’ |
| Cadherin 3 human | 5’ -ATGACTTCACTGTGCGGAATGGCGAGACA- 3’ |

shRNA sequences used for lentiviral gene depletion

**Table S3.**

| Antibody | Clone | Source | Host | Application |
|---|---|---|---|---|
| P-Cadherin | 6A9 | Invitrogen (MA1-2003) | Mouse mAb | WB: 1:1,000 |
| | | Thermofisher (132000z) | Rat mAb | IF: 1:200 |
| COL17 | | Invitrogen (PA5-101001) | Rabbit pAb | WB: 1:1,000 / IF: 1:200 |
| Keratin 14 | Poly19053 | BioLegend (905301) | Rabbit pAb | IF: 1:1,000 |
| Laminin $\alpha$3 mouse | | Dr. Takako Sasaki (1110+) | Rabbit | IF: 1:500 |
| Laminin $\gamma$2 | D4B5 | Millipore (MAB19562-AF647) | Mouse mAb | IF: 1:150 |
| $\beta$-Catenin | E5 | Santa Cruz (sc-7963) | Mouse mAb | WB: 1:500 / ChIP: 2 μg |
| Phospho-FAK.Y397 | D20B1 | Cell Signaling (8556) | Rabbit mAb | WB: 1:1,000 |
| FAK | | Cell Signaling (3285) | Rabbit | WB: 1:1,000 |
| Phospho-Src.Y416 | D49G4 | Cell Signaling (6943) | Rabbit mAb | WB: 1:1,000 |
| Src | | R&D System (AF3389) | Goat pAb | WB: 0.5 μg/ml |
| TCF4 | | Millipore (CS204338) | Mouse mAb | ChIP: 2 μg |
| $\beta$-actin | AC15 | Sigma-Aldrich (A1978) | Mouse mAb | WB: 1:1,000 |

Primary Antibody list

**Table S4.**

| Antibody | Source | Host | Application |
| --- | --- | --- | --- |
| Anti-rabbit IgG, HRP-linked antibody | Cell Signaling Technology (7074S) | Goat | WB: 1:2,000 |
| Anti-mouse IgG, HRP-linked antibody | Cell Signaling Technology (7076S) | Horse | WB: 1:2,000 |
| Alexa Fluor 488 goat anti-rabbit IgG (H+L) | Invitrogen (A11008) | Goat pAb | IF: 1:500 |
| Alexa Fluor 594 goat anti-rabbit IgG (H+L) | Invitrogen (A11012) | Goat pAb | IF: 1:500 |

Secondary Antibody list

**Table S5.**

| Gene | Primer sequence |
|---|---|
| Laminin $\alpha$3 mouse | Forward: 5' -ACACCTGGGACGTGGATTG- 3' |
| | Reverse: 5' -CTTGCAGGGTGAATGCTTCAT- 3' |
| Laminin $\beta$3 mouse | Forward: 5' -GGCTGCCTCGAAATTACAACA- 3' |
| | Reverse: 5' -ACCCTCCATGTCTTGCCAAAG- 3' |
| Laminin $\gamma$2 mouse | Forward: 5' -CAGACACGGGAGATTGCTACT- 3' |
| | Reverse: 5' -CCACGTTCCCCAAAGGGAT- 3' |
| Cadherin 3 mouse | Forward: 5' -CTGGAGCCGAGCCAAGTTC- 3' |
| | Reverse: 5' -GGAGTGCATCGCATCCTTCC- 3' |
| $\beta$-actin mouse | Forward: 5' -GGCTGTATTCCCCTCCATCG- 3' |
| | Reverse: 5' -CCAGTTGGTAACAATGCCATGT- 3' |

Real-time (RT) PCR primers

**Table S6.**

| Gene | Primer sequence |
|---|---|
| Laminin $\alpha 3$ promoter mouse | Forward: 5' -GGCTGGCCTCAAACTCAAGA- 3' |
| | Reverse: 5' -GCTCCCGCTACTATAAGGGC- 3' |
| Laminin $\beta 3$ promoter mouse | Forward: 5' -TTTGGCTTGGGTGCTTTTGG- 3' |
| | Reverse: 5' -CAATGGCAAGAACCTGCGAG- 3' |
| Laminin $\gamma 2$ promoter mouse | Forward: 5' -AGAGATGTCGGCTTGTCTGC- 3' |
| | Reverse: 5' -GTGACCAGTGTAGGCTGACC- 3' |
| SP5 mouse | Forward: 5' -GGGTCTCCAGGCGGCAAG- 3' |
| | Reverse: 5' -AGCGAAAGCAAATCCTTTGAATCC- 3' |

Primers used for Chromatin IP qPCR
